# Proteobacteria with chemosynthetic potential are highly enriched in the gills of *Hypoplectrus* reef fishes

**DOI:** 10.1101/2025.07.01.662686

**Authors:** Sabrin Abdelghany, Martin Helmkampf, Matthew S. Schechter, Iva A. Veseli, Matthieu Leray, A. Murat Eren, Oscar Puebla

**Affiliations:** Leibniz Centre for Tropical Marine Research (ZMT), Bremen, Germany; Institute for Chemistry and Biology of the Marine Environment (ICBM), School of Mathematics and Science, Carl von Ossietzky Universität Oldenburg, Oldenburg, Germany; National Institute of Oceanography and Fisheries (NIOF), Cairo 11516, Egypt; Committee on Microbiology, University of Chicago, Chicago, IL 60637, USA; Helmholtz Institute for Functional Marine Biodiversity at Oldenburg, 26129 Oldenburg, Germany; Smithsonian Tropical Research Institute (STRI), Panama City, Panama, Republic of Panama; Alfred Wegener Institute, Helmholtz Centre for Polar and Marine Research, 27570 Bremerhaven, Germany; Max Planck Institute for Marine Microbiology, 28359 Bremen, Germany

**Author notes:** (S. Abdelghany), (O. Puebla).

## Abstract

A variety of marine invertebrates are known to form associations with chemosynthetic bacteria, but to the best of our knowledge this has not been documented in fishes. Here, we apply genome-resolved metagenomics to the hamlets (*Hypoplectrus* spp), a model system for the study of speciation in the sea. The analysis of 304 gill samples from 12 hamlet species collected at six locations over 13 years revealed a stark contrast between the gill microbiota and ambient water microbial communities. One novel lineage in the Burkholderiaceae-B family was particularly prevalent across host species, sampling locations and years. Its genome encoded highly complete metabolic modules for carbon fixation and sulfur oxidation, indicating chemosynthetic potential. Its pangenome revealed large-scale geographic structure (western Caribbean, eastern Caribbean and Gulf of Mexico), paralleling the phylogenomic pattern observed in the hamlet radiation. Our survey also identified genomes of multiple novel gill-associated lineages related to known fish gill pathogens, fish gut microbes and free-living seawater taxa. These lineages harbor diverse metabolic modules, involved notably in nitrogen cycling, antibiotic production and biofilm formation, revealing a highly dynamic microbial ecosystem. Overall, our findings suggest complex host-microbe and microbe-microbe eco-evolutionary interactions that may influence fish physiology, homeostasis and immune response.

## INTRODUCTION

The ability of some bacteria to fix carbon dioxide and produce energy through the oxidation of inorganic compounds was discovered more than a century ago. This biological process, now referred to as chemolithoautotrophy, is a form of chemosynthesis—as opposed to photosynthesis whereby energy is obtained from the sun. The discovery of hydrothermal vents in 1977 led to the realization that chemosynthetic bacteria can form symbiotic associations with marine invertebrates. Such associations were thereafter identified in a variety of other habitats including cold seeps, whale and wood falls, continental slope sediments, as well as shallow-water sediments associated with coral reefs, seagrasses, mangroves and salt marshes [1, 2]. They involve a diversity of marine taxa including annelids, molluscs, arthropods, nematodes, flatworms, sponges and ciliates. The bacteria can live at the surface or within their host, at the surface or within cells, in different locations including the gut, mantle cavity, specialized structures such as the trophosome in worms, or the gills in molluscs and arthropods [1, 2].

To the best of our knowledge, associations between chemosynthetic bacteria and fishes have not been documented, but if such associations were to occur the gills would be a prime candidate location. The gills constitute the major interface between the body and its environment through which gas exchange occurs. They also contribute to eliminate nitrogenous waste [3, 4] and maintain homeostasis through the regulation of osmotic pressure, pH, and temperature in some cases [4, 5, 6]. Furthermore, the gills constitute a mucosal barrier that contributes to defense against pathogens and maintenance of the fish immune response [7, 8]. Understanding the physiology of gills is therefore critical to assess and predict the adaptability of fishes to rapidly changing natural habitats and to aquaculture environments.

Fish gills also harbor a community of micro-organisms, the so-called microbiome [9, 10, 11]. Through its interactions with mucosal immunity [12] and its potential contribution to ammonia detoxification [13, 14], the gill microbiome may play an important role in fish physiology and health. Metabarcoding, the high-throughput sequencing of 16S ribosomal RNA (rRNA) amplicons, revealed a diverse microbiome dominated by the bacterial phyla Proteobacteria, Actinobacteria, Bacteroidota and Firmicutes [9, 10, 11, 15, 16, 17]. Nevertheless, while some fish gill-associated bacteria have been successfully cultivated and characterized [18, 19], the vast majority of microorganisms that are detected in metabarcoding surveys are missing in culture and their functional role is poorly understood [20, 21]. By enabling the recovery of microbial genomes directly from environmental samples [22], shotgun metagenomics yields new opportunities to characterize the genomes of gill-associated bacteria without cultivation, offering an improved understanding of the fish gill microbiome [23]. Yet a large fraction of studies on the fish microbiome focus on the gut [17], and the studies that target fish gills with shotgun metagenomics tend to focus on specific pathogens [24, 25, 26].

Here, we apply genome-resolved metagenomics to characterize the taxonomic composition and functional potential of the gill microbiome at the scale of an entire radiation, the hamlets (*Hypoplectrus* spp). Hamlets are a group of fishes found throughout the greater Caribbean, primarily associated with coral reefs and feeding on small fishes and invertebrates. The group includes 19+ species [27, 28] that are ecologically very similar [29, 30], highly sympatric (up to nine species on a single reef), and differ essentially in terms of color pattern and mate choice. They are extremely close genetically and form a very shallow radiation that appears to be exceptionally recent and rapid, making them a rare model system for the study of speciation and radiation in the marine environment [31, 32, 33, 34]. In this context, we shotgun-sequenced over 300 hamlet gill tissue samples from 12 species and six locations collected over 13 years (Fig. **1**). This dataset was used to study the host genome [33, 34, 35, 36, 37], but also provides the opportunity to characterize the gill microbiome of the radiation at the metagenomic level, and to investigate how dominant microbial taxa may have co-evolved with their hosts during rapid diversification.

**FIG 1.**
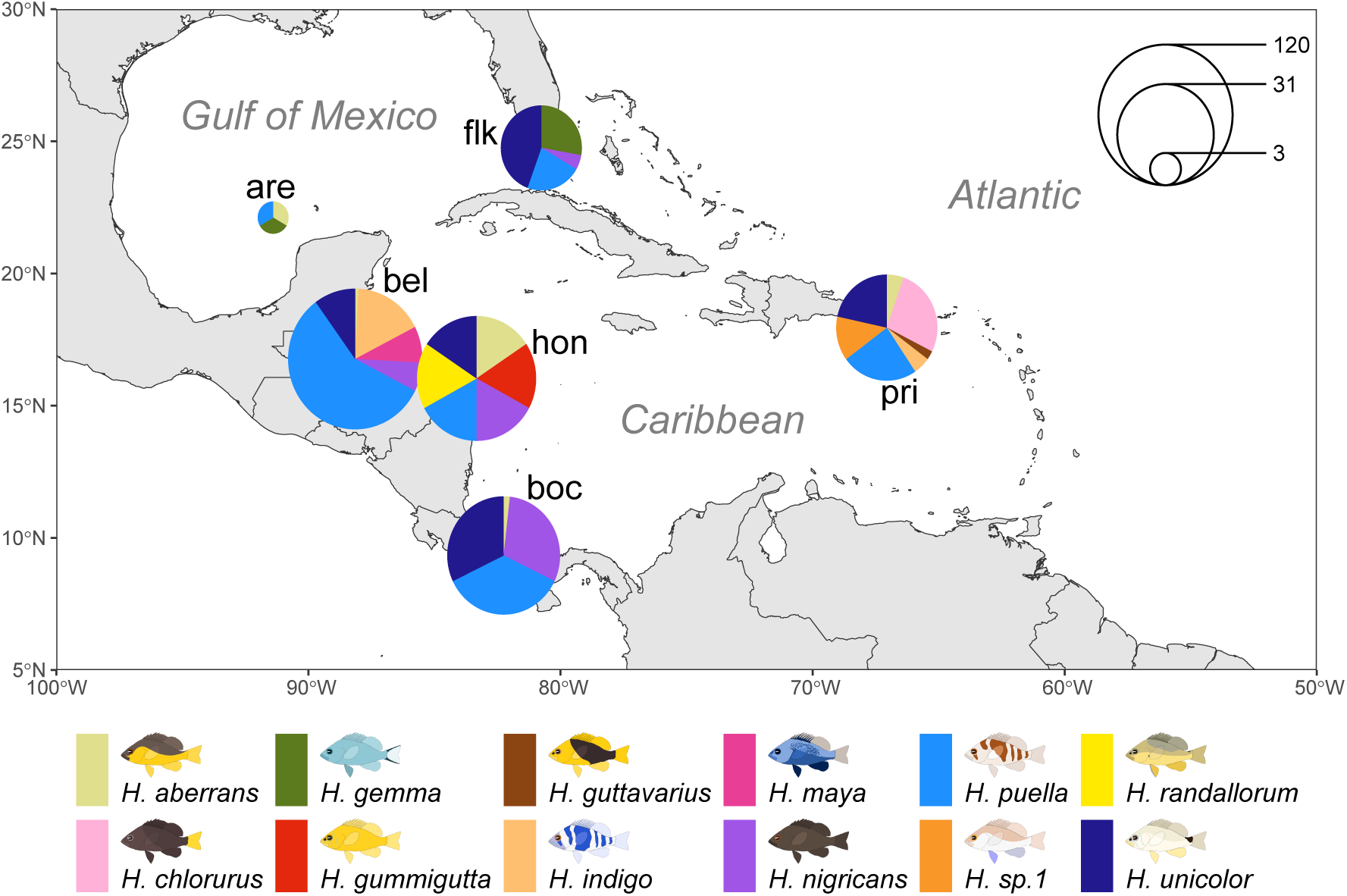
Sampling design. The dataset consisted of 304 shotgun-sequenced gill tissue samples from 12 hamlet species collected at six greater Caribbean locations: Bocas del Toro in Panama (boc), Belize (bel), Puerto Rico (pri), Cayo Arenas in the Gulf of Mexico (are), Honduras (hon) and the Florida Keys (flk).

## RESULTS AND DISCUSSION

### The gill microbiome is diverse and conserved across species, locations and years

We started by characterizing the taxonomic composition of the gill microbiome at the phylum, class and order levels using a genomic read classification approach (Suppl. Figs. 1-4). A total of 1.5 billion quality-filtered reads were retained after removing host sequences, representing 3.9% of the original dataset (37.5 billion reads). Of these, 283 million reads (0.8% of the original dataset) were retained after further filtering out reads classified as animal, plant, cyanobacteria and phi X, as well as unclassified reads. In agreement with metabarcoding studies [9, 10, 11, 15, 16, 17], the results indicated a high prevalence of Proteobacteria (Gamma- and Betaproteobacteria in particular), Actinobacteria, Bacteroidota and Firmicutes in the gills. Nevertheless, the composition of the hamlet gill microbiome differed from other marine and reef fishes at low taxonomic levels (order and below). For example, a high prevalence of *Shewanella*—a genus of Gammaproteobacteria in the order Alteromonadales—was reported in other tropical marine fishes [10, 11, 16], but Alteromonadales was not identified as a dominant order by our read classification analysis. This result is in line with evidence that the microbiome of reef fishes is partitioned among species [38, 39]. Furthermore, we identified protists, fungi, archaea and viruses, providing a taxonomically broader view of the fish gill microbiome diversity than metabarcoding. The occurrence of non-bacterial phyla was not anecdotal, as illustrated by e.g. the high prevalence of Ascomycota reads. We also note the occurrence of Euglenozoa, a group of protists that includes fish gill parasites [40].

The composition of the gill microbiome showed variability among samples, but only 4.7%, 7.0%, 8.8% and 17.9% of this variation was explained by biogeographic region (Gulf of Mexico, western Caribbean and eastern Caribbean), host species, location and sampling year, respectively (Suppl. Table 1). These results point to a gill microbiome that is taxonomically and geographically conserved at the scale of the hamlet radiation. The similarity of the gill microbiome among host species was expected at this taxonomic resolution considering the high genetic [33, 34, 41] and ecological [29, 30] similarity of hamlet species. The most notable variation was among individuals, with some individuals from diverse host species, locations and sampling years showing an atypical community composition. These samples often presented an unusually high proportion of potentially pathogenic taxa such as viruses or bacilli. Diseases are known to have a marked effect on the composition of the gill microbiome [42, 43, 44], suggesting that part of the variation in the composition of the gill microbiome that we observe in the hamlets relates to fish health status and immune response. In this regard the microbial genomes that we present here provide potential for a biomarker-based approach to monitor the health of fish populations by analyzing the composition of their gill microbiomes.

### The gill microbiome is distinct from surrounding seawater

In order to evaluate to what extent the gill microbiome differs from surrounding water, we also analyzed shotgun sequences from reef water collected at one of our sampling locations (Bocas del Toro, Panama) [45, 46]. For this we used the EcoPhylo workflow [47], a novel approach that profiles the distribution of gene family homologs across metagenomic samples, considering the ribosomal protein single-copy core gene *rpL16* as a proxy for the presence of microbial populations.

The results showed little overlap between the gill and seawater microbiomes, with just five taxa out of 93 detected in both environments (Fig. **2**). The dichotomy between the two environments was particularly striking for one Burkholderiaceae-B taxon, which was recovered in 183 out of 197 gill samples but in none of the 19 seawater samples, indicating high prevalence and strong enrichment in the gills compared to seawater. Similar results were obtained with another gene (*rpL13*, Suppl. Fig. 5), indicating that this pattern is not marker-specific. These results stand in stark contrast to previous studies that showed more overlap between gill and seawater microbiomes [10, 11, 48]. This may be due in part to the fact that the metagenomic approach based on the ribosomal protein loci that we used provides greater variability and resolution than metabarcoding based on the 16S rRNA gene [49], which may have contributed to resolving taxa that were lumped with metabarcoding. Furthermore, gill tissue samples were rinsed with sterile water prior to DNA extraction to remove salt from the DMSO buffer in which they were preserved. This may have contributed to reducing contamination from seawater, but on the other hand the drop in osmotic pressure may have burst open microbial cells at the surface of the tissue, biasing our dataset towards endosymbionts. Finally, the hamlets are not filter-feeders, a feeding mode that is expected to result in higher occurrence of seawater material in the gills and therefore a higher similarity between the two microbiomes [50].

**FIG 2.**
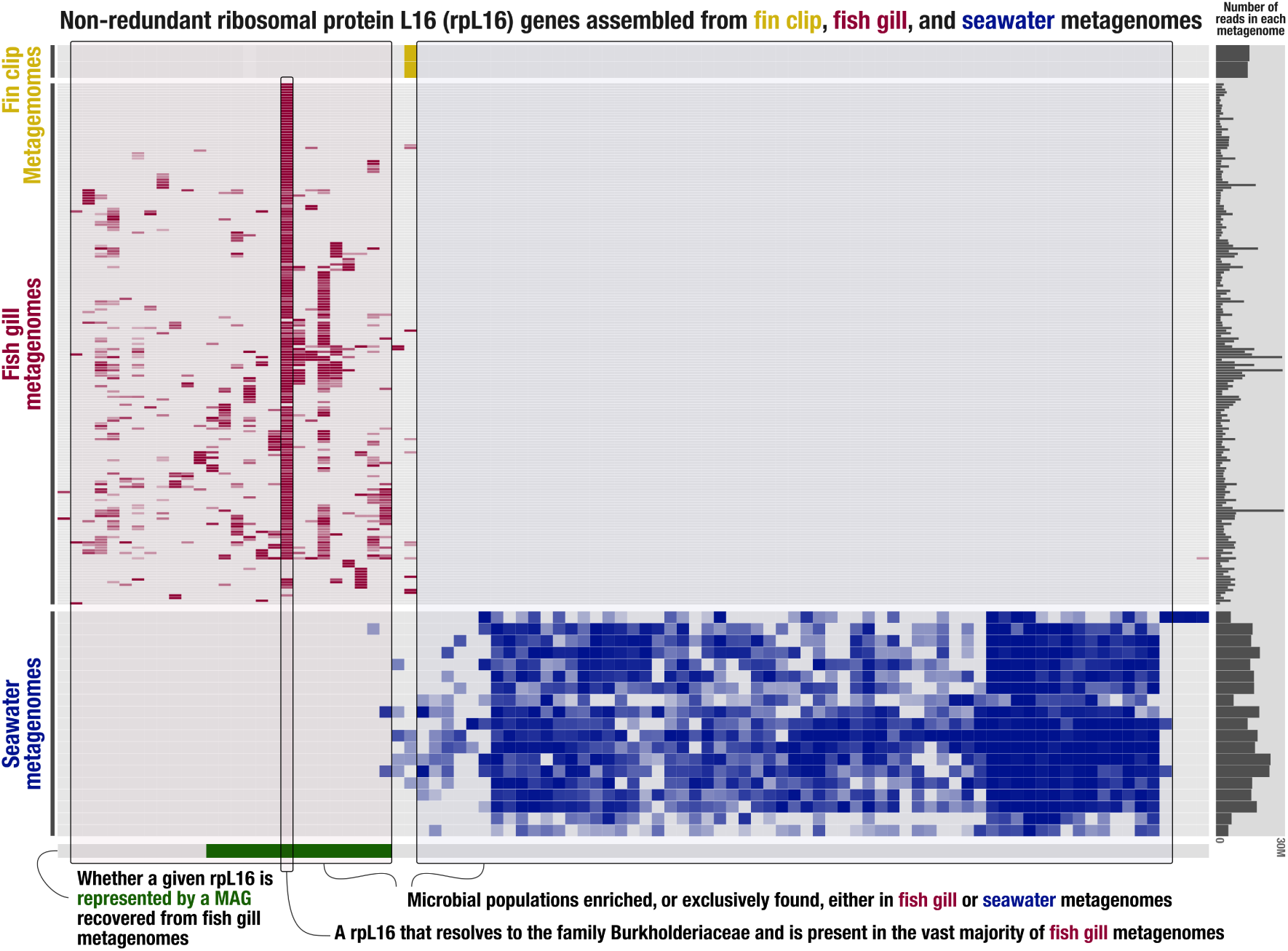
Microbiome community composition in the gills and seawater based on the *rpL16* gene family. The heatmap shows the coverage (0-100%) of the different *rpL16* clades (columns) in the gills versus seawater metagenomes (rows), indicating little overlap in community composition between the gill and seawater microbiomes with just five clades out of 93 detected in both environments. Note the high prevalence of one clade in the gills (and conversely its absence in seawater), which corresponds to a novel taxon in the Burkholderiaceae-B family. Another striking pattern is the lower microbial diversity of the gills compared to seawater, indicating a more specialized microbiome in the gills. The green part of the horizontal bar at the bottom represents the *rpL16* clades that were recovered in gill MAGs, indicating that despite the large sampling and sequencing effort, the bacterial lineages that were recovered as MAGs from the gills represent just 54% of the lineages detected by the *rpL16* marker in the gills.

The EcoPhylo analysis also showed a lower diversity of the gill microbiome compared to coral reef water, which is consistent with previous studies [10, 51] (but see [11]), suggesting a specialized microbiome in the gills. All in all, our analyses point to a specific gill microbiome that is less diverse and distinct from seawater, underscoring the particular selective pressures that characterize this specific habitat. This raises the question as to how the microbiome colonizes the gills. Horizontal transmission is likely since fertilization is external and both eggs and larvae are pelagic in the hamlets. This would imply that the taxa identified in the gills are present in the environment, albeit below detection level in the seawater samples that we analyzed.

### A diversity of MAGs were assembled, most of which belong to novel species

A total of 67 MAGs (Metagenome-Assembled Genomes) were assembled from the dataset (Table 1). The average assembly completeness across all retained MAGs was 81.8% (min. 42.2%, max. 99.3%), and the average contamination was 1.5% (min. 0%, max. 7.6%). The MAGs belonged to three bacterial phyla (Pseudomonadota, Chlamydiota and Bacteroidota), four classes (Gammaproteobacteria, Chlamydiia, Alphaproteobacteria and Bacteroidia), eight orders and ten families according to the Genome Taxonomy Database (GTDB), and the phylogenomic analyses below indicate that they represent 17 distinct bacterial lineages. Following a 95% Average Nucleotide Identity (ANI) cut-off [52, 53, 54], 15 of these 17 lineages would represent new species. Some of these may also represent new genera, and six MAGs in the order Paracaedibacterales could not be assigned to any known family. Furthermore, the EcoPhylo analysis indicates that despite the extensive sampling and sequencing effort, the bacterial lineages that were recovered as MAGs from the gills represent just 54% of the lineages detected by the *rpL16* marker in the gills (Fig. **2**).

**TABLE 1.**
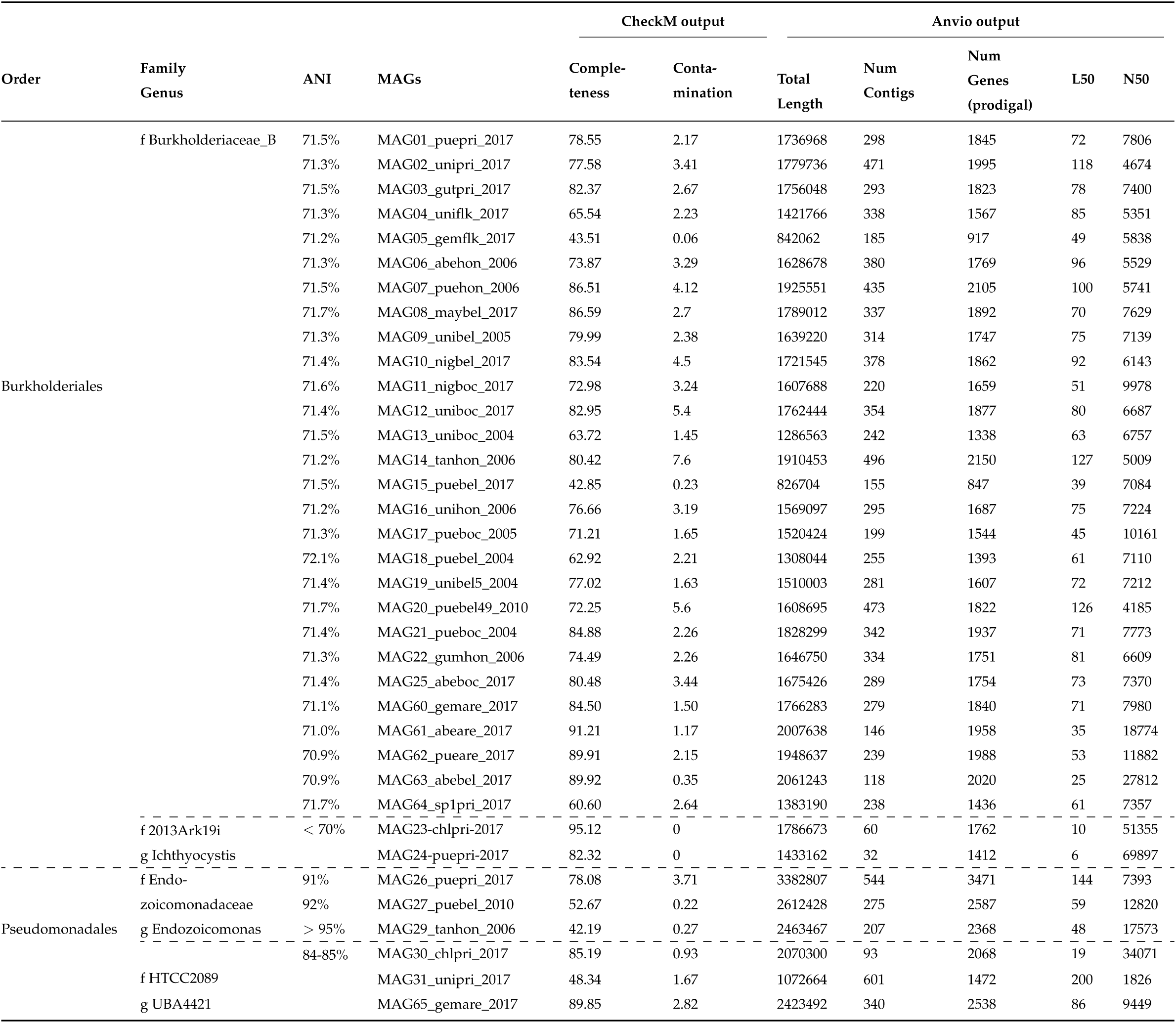
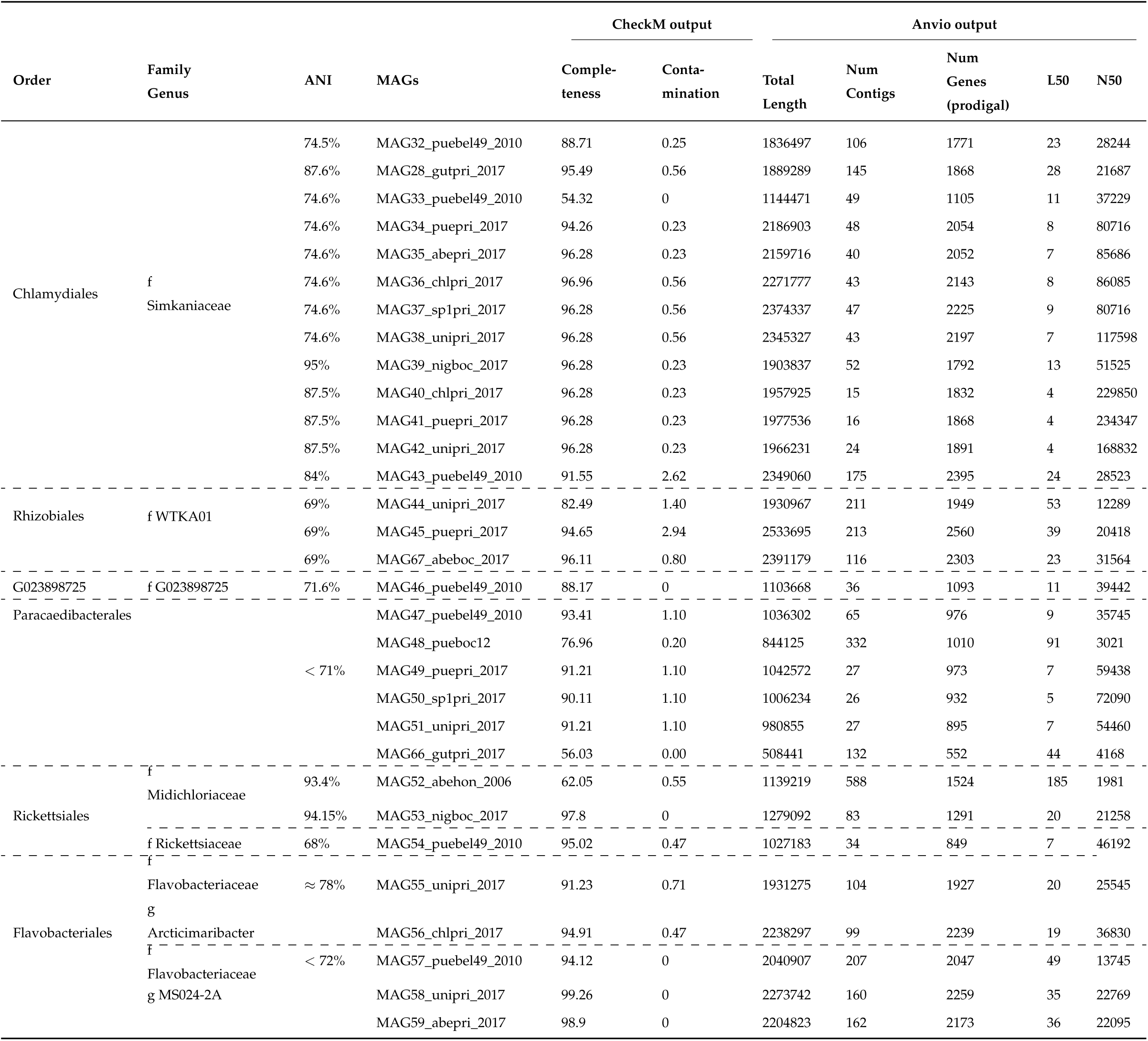
The 67 Metagenome-Assembled Genomes (MAGs) recovered in this study. ANI: Average Nucleotide Identity with the ost closely related species. The host species and location are abbreviated in the MAG names as described in Suppl. Fig. 5 caption.

We focused on the Burkholderiales, Pseudomonadales, Rhizobiales, and Flavobacteriales orders for more in-depth analysis. These orders were selected because they host MAGs encoding a diversity of metabolic modules. They represent 65% of the MAGs and half of the orders that were recovered. The remaining four orders represent just 23 MAGs encoding a limited number of metabolic modules notwithstanding high genome completeness (mean 88.5%, Table 1), suggesting that they might be obligate symbionts or pathogens, which will be the focus of another study.

The results of the phylogenomic analyses for the Burkholderiales, Pseudomonadales, Rhizobiales, and Flavobacteriales are presented in Suppl. Figs. 8-18. Some MAGs were related to fish gill pathogens, notably *Ichthyocystis sparus* and *I. hellenicum* [25] in the Burkholderiales (Suppl. Fig. 8), as well as *Endozoicomonas sp900299555* and *E. elysicola* [55, 24] in the Pseudomonadales (Suppl. Fig 10). The genus *Endozoicomonas* is known for its association with corals, but also includes pathogenic and parasitic lineages of other marine animals [56]. Other MAGs from this genus were identified as close relatives of *E. atrinae*, which was isolated from the gut of the comb pen shell (*Atrina pectinata*) [57]. Yet other MAGs related to predominantly free-living seawater genera, notably *UBA4421* in the Pseudomonadales (Suppl. Fig 12) as well as *MS024-2A* and *Arcticimaribacter* in the Flavobacteriales (Suppl. Fig. 16). Finally, some MAGs were found to be related to *WTKA01 sp011524605* in the Rhizobiales, which is associated with coral reef macroalgae biofilm (Suppl. Fig. 14). These results indicate that the microbes found in the gills are phylogenetically related to bacteria that are adapted to a variety of ecological contexts.

### The most prevalent Burkholderiaceae-B taxon co-evolved with the host

The Burkholderiaceae-B taxon that was most prevalent in the gills according to the EcoPhylo analysis (Fig. **2**) represented 28 of the 67 MAGs across ten host species, six sampling locations and five sampling years between 2004 and 2017 (Table 1). It was followed by two species in the Chlamydiales and Paracaedibacterales orders that were recovered as MAGs just six times each, confirming the high prevalence of the Burkholderiaceae-B taxon. Considering that this taxon is novel, its high prevalence in the gills but absence in seawater, and the fact that the gill dataset encompasses different sampling expeditions, DNA extraction batches and sequencing runs over several years, it is unlikely that it represents a contamination. Furthermore, we analyzed 16 fin clips from different hosts and locations as controls and recovered no MAG and a single *rpL16* clade (*Vibrio*) from these samples (Fig. **2**), ruling out a systematic contamination of the hamlet samples by the Burkholderiaceae-B taxon.

Phylogenomic analysis indicated that the Burkholderiaceae-B MAGs form a deeply branching clade in the family relative to known species (Fig. **3**). The ANI values to the most closely related species ranged between 70.9% and 72.1% (Table 1), and the identity of the most closely related species varied from MAG to MAG. Furthermore, these ANI values are based on the sections of the genome that align, and these represented a small proportion of the genome (≤ 20%), indicating that the Burkholderiaceae-B MAGs are distantly related to any known species in the family.

**FIG 3.**
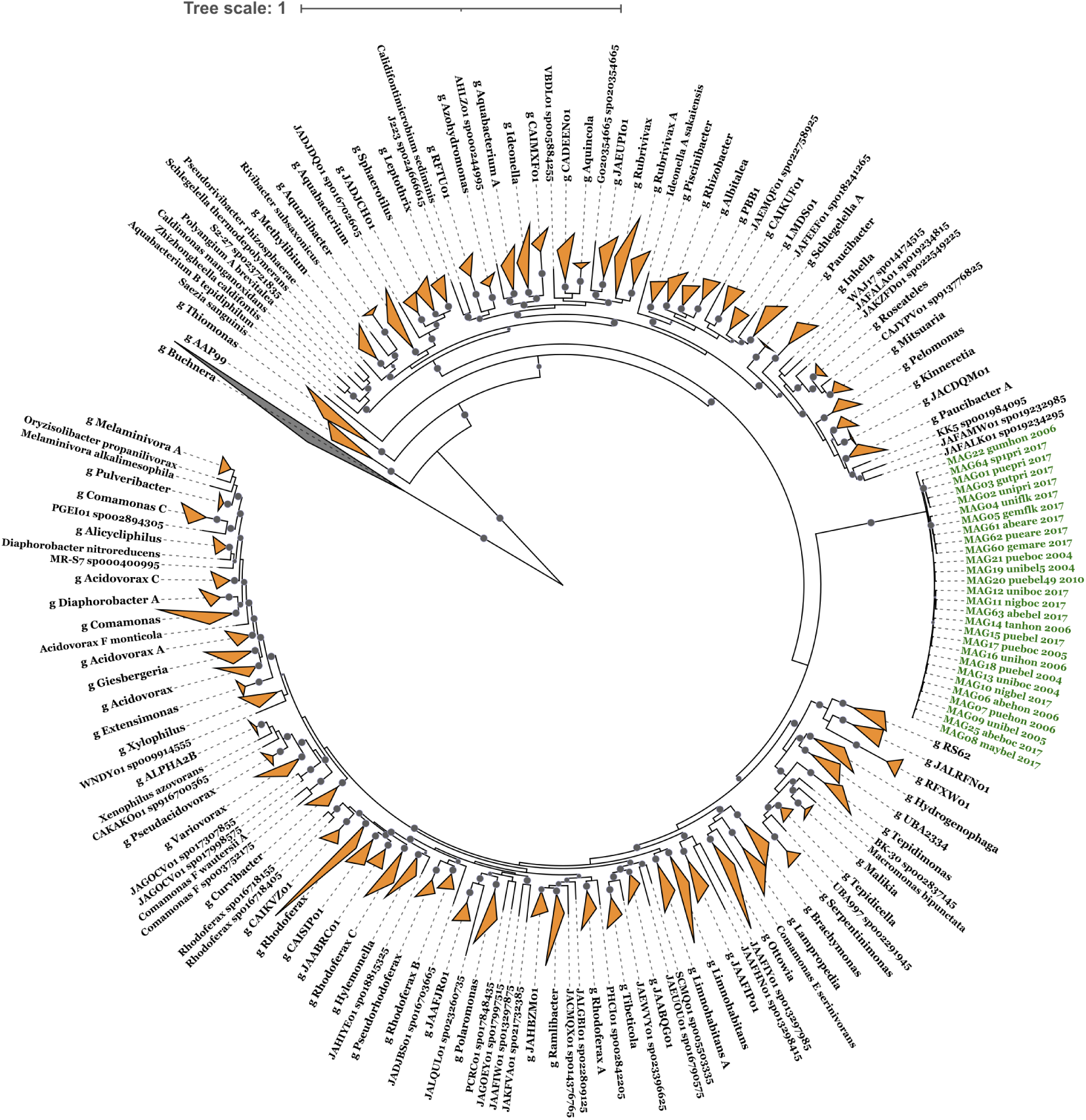
Phylogenomic placement of the MAGs from the most prevalent taxon in the Burkholderiaceae-B family. These 28 MAGs (in green) form a distinct clade that is ∼ 70% diverged from the most closely related species based on average nucleotide identity. Genus names are indicated by a leading "g" where genus nodes were collapsed. Triangles size represent the number of genomes in these collapsed nodes. The tree is rooted with the genus *Buchnera* (in black). Bootstrap values represented by gray circles, only bootstrap >95 are shown.

**FIG 4.**
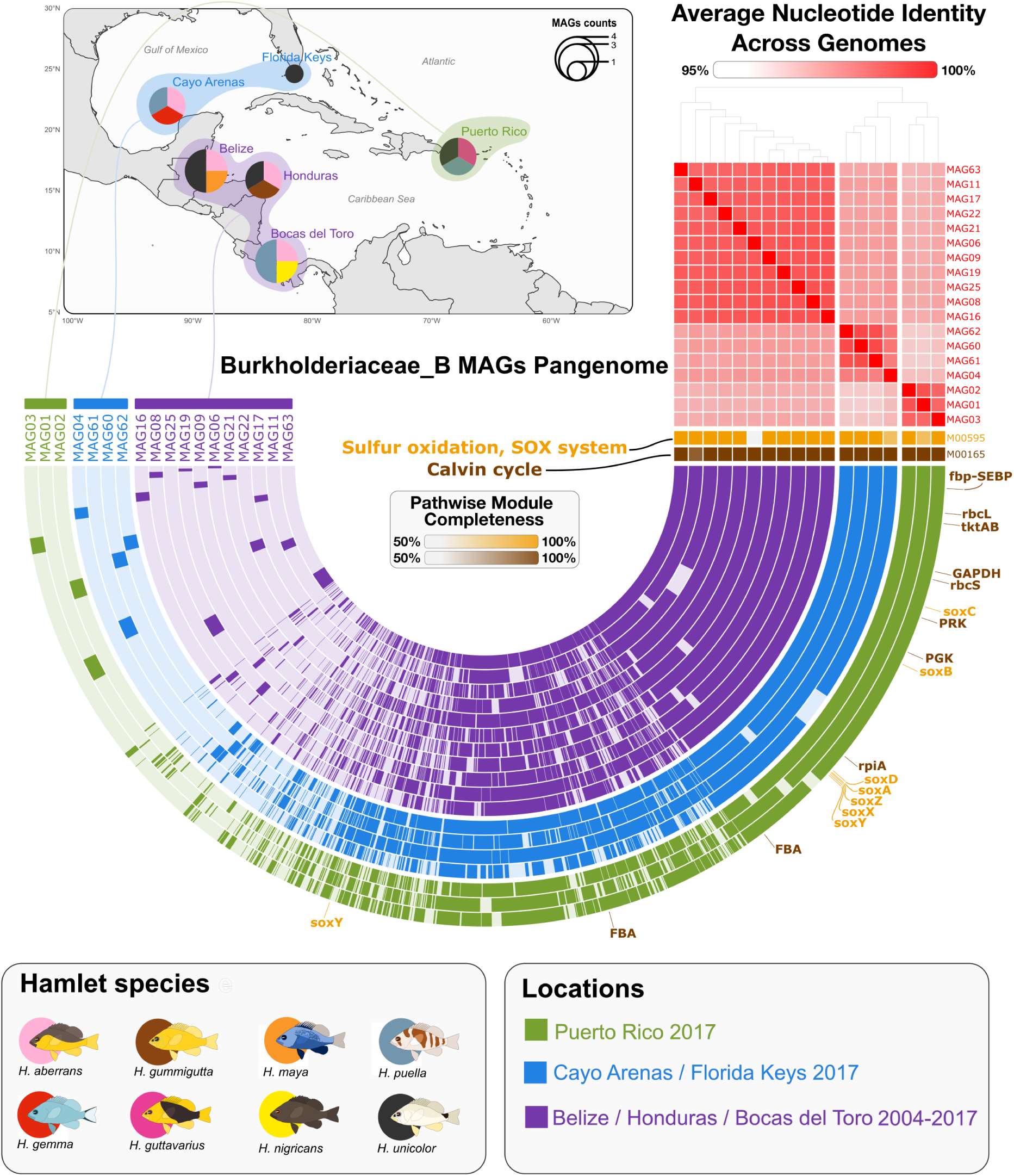
**Pangenome of the most prevalent Burkholderiaceae-B taxon**. Each layer represents one MAG. The heatmap in the top-right shows average nucleotide identity among all genomes. A total of 4478 gene clusters were identified, including 196 core gene clusters (recovered in all genomes), 1768 accessory gene clusters (recovered in only part of the genomes), and 2227 singleton gene clusters (recovered in just one genome). A heatmap of pathwise module completeness scores for the Sulfur oxidation module (yellow) and the Calvin cycle module (brown) is shown below the ANI heatmap. Gene clusters containing the genes belonging to these modules are indicated by gene name labels around the outside of the pangenome. For the Calvin cycle, these genes include fbp-SEBP: bifunctional fructose-1,6/sedoheptulose-1,7-bisphosphatase; rbcL: ribulose-bisphosphate carboxylase, large chain; rbcS: ribulose-bisphosphate carboxylase; small chain; tktAB: transketolase; RPE: ribulose-phosphate 3-epimerase; GAPDH: glyceraldehyde-3-phosphate dehydrogenase; PRK: phosphoribulokinase; PGK: phosphoglycerate kinase; rpiA: ribose-5-phosphate isomerase; FBA: fructose-bisphosphate aldolase; and TPI: triosephosphate isomerase. Note that enzymes TPI and RPE are not included in the version of KEGG module used in our analysis and were manually identified. For the SOX system, these genes include the sulfur-oxidizing proteins soxABXYZ and the S-disulfanyl-L-cysteine oxidoreductases soxCD.

The 28 Burkholderiaceae-B MAGs shared 95.7–98.3% ANI and had a mean alignment fraction of 73.8%, indicating that they would belong to the same species following a 95% ANI and a 50% alignment fraction threshold [52, 53, 54, 58].

The high number of Burkholderiaceae-B MAGs allowed us to generate a pangenome for the group (Fig. **4**). It revealed broad-scale geographic structure, with samples from the Gulf of Mexico, eastern Caribbean and western Caribbean grouping together regardless of host species and collection year. This pattern was confirmed by phylogenomic analysis (Suppl. Fig. 7). It reflects the phylogenomic structure observed in the hamlet radiation, which is poorly structured with regard to host species but shows a biogeographic signal among the same three regions [34]. This indicates possible co-evolution between the Burkholderiaceae-B taxon and its host, albeit at the biogeographic not taxonomic level. These results resonate with what has been observed in lucinid clams, where some chemosynthetic symbionts show low host specificity but geographic partitioning [59].

### The most prevalent Burkholderiaceae-B taxon has chemosynthetic potential

Functional analysis of the 28 Burkholderiaceae-B MAGs identified the complete set of Calvin cycle enzymes for carbon fixation (Fig. 4, Suppl. Fig. 19, Suppl. Table 2), indicating autotrophic potential. This is in line with the observation that many members of the Burkholderiaceae host a complete Calvin cycle [60]. Genes involved in the dark phase of the Crassulacean Acid Metabolism (CAM) pathway were also present, but in bacteria these genes might contribute to other functions than carbon fixation as they do in plants. Genes encoding the full Sox complex (soxABCDXYZ) for aerobic thiosulfate oxidation were also recovered. The Calvin cycle and thiosulfate oxidation pathways were recovered complete not only in genomes co-assembled from multiple hosts, but also in genomes assembled from a single host, indicating that they co-occur in the same host. Altogether, the Calvin cycle and thiosulfate oxidation pathways provide potential for chemosynthesis, chemolithoautotrophy more specifically. Chemolithoautotrophic thiosulfate oxidizers have been reported in the Burkholderiales, albeit in different ecological contexts. Notably, they have been identified in the soil rhizosphere [61], but these were facultative chemolithoautotrophs. In our case the lack of a complete glycolysis pathway suggests that chemolithoautotrophy is obligate.

Sulfide:quinone oxidoreductase, which is essential to link reduced sulfur compounds to the quinone pool, was consistently present across MAGs. Furthermore, multiple NADH:quinone oxidoreductase subunits, the full bc1 complex and cytochrome c oxidase were also recovered. This suggests that reduced quinones may transfer electrons to cytochrome c, which are then passed to oxygen via a terminal cytochrome c oxidase, indicating capacity for aerobic respiration. Additional bd-type oxidases were also recovered, suggesting ability to cope with microaerobic conditions. Enzymes related to taurine (taurine dehydrogenase) and glycine betaine (glycine betaine/proline uptake), two universal osmolytes, were also present. Beyond their functions as osmolytes, taurine may contribute to sulfur cycling in the gills [62] and glycine betaine has been suggested as another possible source of carbon in symbiotic interactions [63]. The three subunits of aerobic carbon monoxide dehydrogenase were also found, which resonates with the fact that some Sox symbionts can use carbon monoxide as a source of energy [63, 64].

The high prevalence of the Burkholderiaceae-B taxon suggests that it is a predominant constituent of the hamlet core gill microbiome. In this regard it may also be important for niche occupation, contributing to stabilize the gill microbiome and limit colonization by pathogens. This hypothesis is consistent with the occurrence of components of the CRISPR-Cas immune system in its genome, which is essential for defense against phage invasions [65] and has been suggested to enhance the resilience of sponge microbiomes [66]. More generally, the Burkholderiaceae-B taxon exhibited diverse metabolic capacities, encoding 56 metabolic modules with high completeness (≥ 75%), including 12 amino acids synthesis pathways (Suppl. Fig. 19) as well as a variety of ATP-binding cassette (ABC) transporters, cofactors and vitamins.

Associations between sulfur-metabolizing bacteria and gills have been reported before, but in invertebrates and often in extreme environments such as deep-sea hydrothermal vents or sediments that are rich in reduced sulfur compounds [64, 67, 2]. While the hamlets live in shallow-water coral environments that are not particularly extreme in terms of environmental conditions, the gills may nonetheless provide a specific microhabitat that is enriched in CO_2_ and other compounds generated by the respiration and excretion of the host, which may be conducive to chemosynthesis. All in all, our metagenomic data indicate metabolic potential for chemosynthesis in the most prevalent species in the hamlet gills. Considering that the hamlets are not particularly specialized ecologically, we speculate that such associations may be widespread in fishes, with possible implications for their metabolism and physiology.

### The gill microbiome harbors a variety of metabolic functions

The results of the functional analyses for the Burkholderiales, Pseudomonadales, Rhizobiales, and Flavobacteriales are presented in Suppl. Figs. 20-25. Metabolic modules for carbon fixation were also identified in the Rhizobiales, which here again is in line with the observation that many members of this order host a complete Calvin cycle [60]. We also recovered a number of modules involved in nitrogen cycling. These include nitrate assimilation and dissimilatory nitrate reduction into ammonia (both in *Endozoicomonas*). The nitrogenous waste of the host is excreted as ammonia in the gills, and experimental evidence indicates that the gill microbiome can contribute to detoxify this ammonia through ammonia oxidation and denitrification [13]. We did not identify modules that relate to these specific functions in the MAGs that we functionally analyzed, but ammonium transporters were nonetheless detected in almost half of the MAGs, suggesting active ammonia transport into cells [68]. The nitrogen regulatory protein GlnK, a member of the PII family, was found in 22 of 28 MAGs from the Burkholderiaceae-B lineage and in three additional MAGs. GlnK regulates ammonium assimilation by interacting with the ammonium transporter AmtB, a process modulated by uridylylation via the GlnD enzyme [68, 69]. Besides, we identified genes encoding for the transport of nitrate, nitrite in *Endozoicomonas*, and branched-chain amino acids across the cell membranes of various MAGs, potentially facilitating effective nitrogen utilization and recycling in the gills (Suppl. Table 3).

We also identified several COG20 functional categories associated with colonization and interactions between the host and microbes across pathogenic, commensal, and mutualistic relationships. These include eukaryotic-like proteins such as ankyrin repeats and tetratricopeptide repeats, as well as membrane proteins involved in the export of O-antigen and teichoic acid. Several MAGs also encoded genes for type IV pili that provide potential for biofilm formation and host cell adherence [70], as well as flagella construction and chemotaxis signaling (Burkholderiaceae-B MAGs lineage not included), indicating motility potential (Suppl. Table 4).

Our results further highlight a complex web of potential nutritional interactions within the gill microbiome, notably through the biosynthesis of a diverse array of B vitamins. These include riboflavin (B2), folic acid (B9), pantothenic acid (B5), pyridoxamine (B6), niacin (B3), and cobalamin (B12) across various MAGs. The ability of different MAGs to produce essential vitamins suggests possible beneficial relationships both between microbes and between the microbiome and the host, as suggested for Pseudomonadales in octocorals [71] and gammaproteobacterial symbionts in deep-sea bivalves [72]. Beyond vitamins, the gill MAGs harbored modules for the production of diverse metabolites. For example ectoine (in *Endozoicomonas*), an osmolyte that has been suggested to contribute osmoregulation in marine fishes [73]. Notably, modules for isoprenoid and dTDP-L-rhamnose biosynthesis were widespread in the MAGs. These pathways are linked to signaling, virulence, and symbiosis [74, 75, 76, 77].

## Conclusion

To the best of our knowledge, the metagenomic results presented here constitute the first line of evidence suggesting that fishes may host chemosynthetic bacteria in their gills. Experimental work is required to determine the location of the Burkholderiaceae-B taxon in the hamlet gills, asses the extent to which its chemosynthetic potential is realized *in vivo*, and if so whether it relates to the physiology of the host or other gill microbes. Nevertheless, while our metagenomic study constitutes an advance in the characterization of the fish gill microbiome, what it really highlights is how little is known about its diversity and potential host-microbe and microbe-microbe interactions. The high proportion of novel taxa identified indicates that we are still at an exploration and discovery phase. In this regard more sequencing across a broader diversity of fishes and habitats is required. More generally it is time for a research program aimed at disentangling the complex web of interactions that take place in the fish gill microbiome, in the spirit of van Kessel *et al*. [13]. The metagenomes described here lay a foundation for this work by allowing to develop species-specific markers that can be used for targeted research.

## MATERIALS AND METHODS

### Sample collection and processing

This study includes a total of 304 shotgun-sequenced gill tissue samples from 12 hamlet species collected between 2004 and 2017 at six locations across the greater Caribbean (Fig. **1**, Suppl. Fig. 6, Suppl. Table 5). This represents a total of 38 host species/location/collection year combinations, with a mean sample size of eight samples per host species/location/collection year (min. one, max. 49). Furthermore, we analyzed 16 fin clip samples from different hosts and locations. The objective was not to characterize the skin microbiome—swabs would have been more appropriate for that—but to process and analyze another type of hamlet tissue samples in the same way as the gills, serving as controls. These genomes were originally generated for studies on the host [35, 36, 33, 37, 34]. Briefly, individuals were collected on scuba using either microspears or hook-and-line. Gill or fin tissue samples were dissected, preserved in salt-saturated DMSO and stored at 4 °C. Bulk DNA was extracted from about 2 mm^2^ of tissue using Qiagen MagAttract High Molecular Weight kits, without any particular step for microbial DNA enrichment or host DNA depletion. Extracted DNA was shotgun-sequenced on an Illumina platform (2 × 151 bp) at a coverage of approximately 20x with respect to the host genome. Host reads were identified by mapping them to the host reference genome with bowtie2 [78] and filtered (note that the host genome assembly procedure included a microbial DNA decontamination step [35]).

### Taxonomic composition of the gill microbiome

We started by characterizing the taxonomic composition of the gill microbiome using the read classification tool Kraken2 v2.0.7-beta [79]. The output from Kraken2 was further processed with Bracken v2.9 [80], which uses a Bayesian approach to re-estimate abundances after classification by Kraken. Reads classified as animal, plant, and Cyanobacteria phyla were filtered out, as well as reads assigned to the phi X phage that was used as control in the sequencing runs. The taxonomic composition of the gill microbiome was analyzed at the phylum, class and order levels and displayed with ggplot2 [81] v3.4.3 in R. Differences in gill microbiome composition were tested using Permutational Multivariate Analysis of Variance (PERMANOVA) with the adonis2 function in vegan [82], computed with 999 permutations. Abundance data were aggregated at the phylum level to assess the effects of biogeographic region, geographic location, host species, and sampling year.

### Metagenome assembly

The entire metagenome assembly workflow, from sequence quality control to binning, was conducted on the anvi’o 7.1 platform following the snakemake workflow (https://merenlab.org/2018/07/09/anvio-snakemake-workflows/). The assessment of raw sequence quality was performed with the iu-filter-quality-minoche script [83]. Subsequently, sequences were assembled into contigs with Megahit [84], with a minimum contig length of 1000 bp. For the 31 host species/locations/collection year for which we had more than one sample, metagenomes were co-assembled per host species, location and collection year. Gene prediction was conducted with Prodigal v2.6.3 [85] and single-copy genes were identified with HMMER v3.2.1 [86]. Quality-filtered reads were mapped to the assembled contigs with Bowtie v2.3.5 [78] and SAMtools v1.10 [87]. The contigs were then binned into Metagenome-Assembled Genomes (MAGs) with anvi-cluster-contigs using the unsupervised binning tool CONCOCT [88]. A subsequent supervised binning procedure was executed through the anvi-interactive interface of anvi’o v7.1. Specifically, the MAGs were inspected and refined using anvi-refine. For the seven host species/location/collection year for which we had only one sample (Suppl. Fig. 6), MAGs were binned from single assemblies with MetaBAT2 [89] after assembling contigs with Megahit. Only bins with a completion rate of at least 40% and contamination levels below 10%, as assessed by CheckM [90], were retained for further analysis.

### Gills versus seawater

The anvi’o EcoPhylo workflow (https://anvio.org/help/main/workflows/ ecophylo/) was used to profile the distribution of MAGs recovered from gill samples in metagenomes from gills, fin clips and seawater. Briefly, this workflow [91] (1) identifies gene family homologs in genome and metagenome assemblies with a user-provided profile Hidden Markov Model [86], (2) clusters open reading frames and selects representative sequences with mmseqs2 [92], (3) leverages the representative nucleotide sequences for metagenomic read recruitment to examine the ecological distribution of the protein family, and (4) calculates a phylogenetic tree with the translated open reading frames to explore the phylogenetic diversity of the protein family [93, 94, 95, 96, 78]. Lastly, the workflow displays an interactive figure to explore the phylogeography of the protein family.

We used the ribosomal protein single-copy core gene *rpL16* (recovered in 59 of the 67 MAGs) as a proxy for the presence of a microbial population. The workflow examined its distribution across gill and fin clip metagenomic data and published metagenomes from seawater collected over coral reefs in Bocas del Toro (Panama) in 2017 [45, 46]. This is one of the locations where the gills considered in this study were collected (in 2004, 2005 and 2017). We adjusted the EcoPhylo workflow with the following parameters: cluster-X-percent-sim-mmseqs –min-seq-id parameter set to 95% and –cov-mode to 1. Finally, we repeated the analysis with *rpL13* (recovered in 57 of the 67 MAGs) to assess the consistency of the results with another marker.

### Phylogenomic analysis

Following interactive refinement in anvi’o, MAGs were exported with anvi-summarize for taxonomic classification. Taxonomic assignments were conducted with the Genome Taxonomy Database Toolkit (GTDB-Tk) v2.3.0 using reference data vr214 [97, 98]. The classify workflow was applied using a collection of 120 bacterial gene markers. This involved placing the MAGs in a reference tree based on the Genome Taxonomy Database (GTDB) with FastANI v1.33 [54] and pplacer [99] to obtain a first approximate phylogenetic placement of the MAGs. Subsequently, the de novo workflow in GTDB-tk [97] was used to infer the phylogenetic relationships between the targeted MAGs and related reference genomes. In this workflow, the infer step used FastTree [95] with the WAG+GAMMA model to build *de novo* trees. The trees are then rooted with a specified outgroup and labeled with the GTDB taxonomy. The anvi-compute-genome-similarity tool (https://anvio.org/m/ anvi-compute-genome-similarity) was used with PyANI [100] to quantify the genomic similarity between reconstructed MAGs and the most closely related genomes.

### Pangenome

We leveraged the high prevalence of one novel taxon of Proteobacteria in the Burkholderiaceae-B family (28 MAGs) to generate a pangenome across host species, sampling locations and sampling years for this taxon. In order to minimize bias due to MAG contamination and low completeness, we only considered MAGs with a completeness >70% and a contamination <3.5%, except for one MAG from the Florida Keys as a representative of this area, resulting in 18 MAGs for this analysis. Functional annotations from different sources (KEGG_Class, COG20_FUNCTION, KOfam, COG20_PATHWAY, KEGG_Module and COG20_CATEGORY) were added to the genome databases. The script anvi-pan-genome was then used with the parameter ’minbit’ set to 0.5 and ’mcl-inflation’ to 10 to identify *de novo* gene families and their distribution across genomes based on amino acid sequence homology determined by BLAST [101]. Subsequently, anvi-display-pan was executed and an ANI heatmap was generated with anvi-compute-genome-similarity. The pangenome visualization was polished with Inkscape (https://inkscape.org/).

### Functional analysis

We functionally annotated the contig databases of all MAGs with KOfam, a customized Hidden Markov Model (HMM) database of Kyoto Encyclopedia of Genes and Genomes (KEGG) orthologs [102], using anvi-run-kegg-kofams (https://anvio.org/m/anvi-run-kegg-kofams). Note that we used a version of KEGG downloaded in September 2023 (for reproducibility, the hash of the KEGG snapshot available via anvi-setup-kegg-kofams was a2b5bde358bb). We used the KEGG annotations with anvi-estimate-metabolism (https://anvio.org/m/anvi-estimate-metabolism) [103] to reconstruct metabolic modules and estimate module completeness, implementing the –only-complete flag. Module completeness was defined by the proportion of complete steps, with a threshold of at least 0.75 to consider a module complete. Additionally, we used the NCBI Clusters of Orthologous Genes (COGs) with anvi-run-ncbi-cogs [104] and the European Bioinformatics Institute (EBI) Pfam database [105]. Heatmaps were generated with ggplot2 in R to visualize the completeness of the metabolic modules.

## DATA AVAILABILITY

This research was conducted on existing data. The raw genomic data are publicly available in the European Nucleotide Archive (accession numbers in Suppl. Table 5). The MAGs will be uploaded in the European Nucleotide Archive (ENA) under the project accession number PRJEB89189.

## ACKNOWLEDGMENTS

This research was funded by a German Academic Exchange Service (DAAD) scholarship to Sabrin Abdelghany (57403037) through the German-Egyptian Research Long-Term Scholarship (GERLS). Data analyses were performed on the high-performance computing cluster ROSA (University of Oldenburg, Germany), funded by the DFG through its Major Research Instrumentation Programme (INST 184/225-1 FUGG) and the Ministry of Science and Culture (MWK) of the state of Lower Saxony. We thank Laetitia Wilkins for initial discussions, Nicole Dubilier, Isidora Morel and Véronique Helfer for feedback on the manuscript, as well as Kosmas Hench for technical support.

## AUTHOR CONTRIBUTIONS

1. S. Abdelghany conducted all the analyses with input and support from all co-authors. S. Abdelghany wrote the first draft of the manuscript and all co-authors contributed to the writing of the final manuscript. All co-authors contributed to the figures. O. Puebla designed the study with input from all co-authors.

## MATERIALS & CORRESPONDENCE

Correspondence and material requests should be addressed to S.A. and/or O.P.

## Supplemental material

**Supplementary Table 1.**
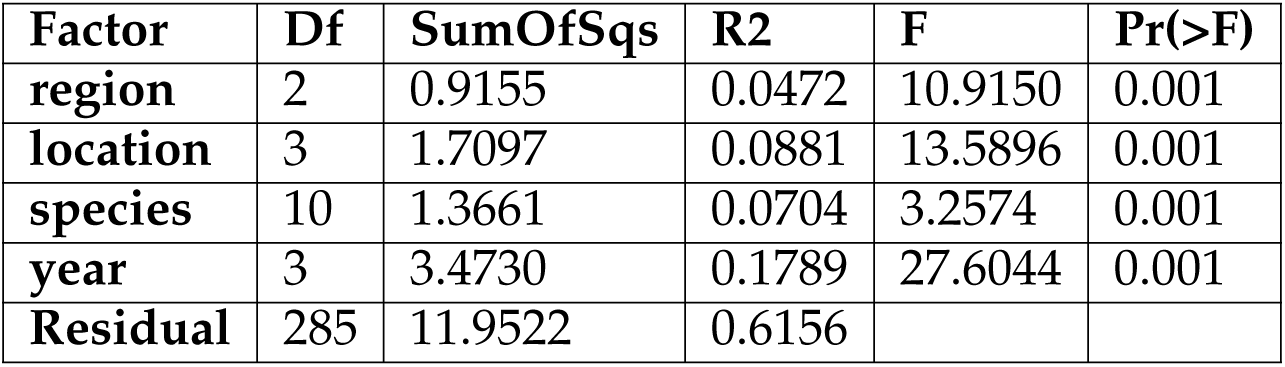
PERMANOVA results for gill microbiome community composition. Results of the adonis2 PERMANOVA assessing the effects of biogegraphic region western Caribbean, eastern Caribbean and Gulf of Mexico), sampling location, host species, and sampling year on gill microbiome composition based on read classification at the phylum level. The table reports the degrees of freedom (Df), sums of squares (SumOfSqs), proportion of variance explained (R^2^), F-statistic (F), and associated p-values (Pr(>F)) based on 999 permutations of a Bray-Curtis dissimilarity matrix.

**Suppl. Figure 1.**
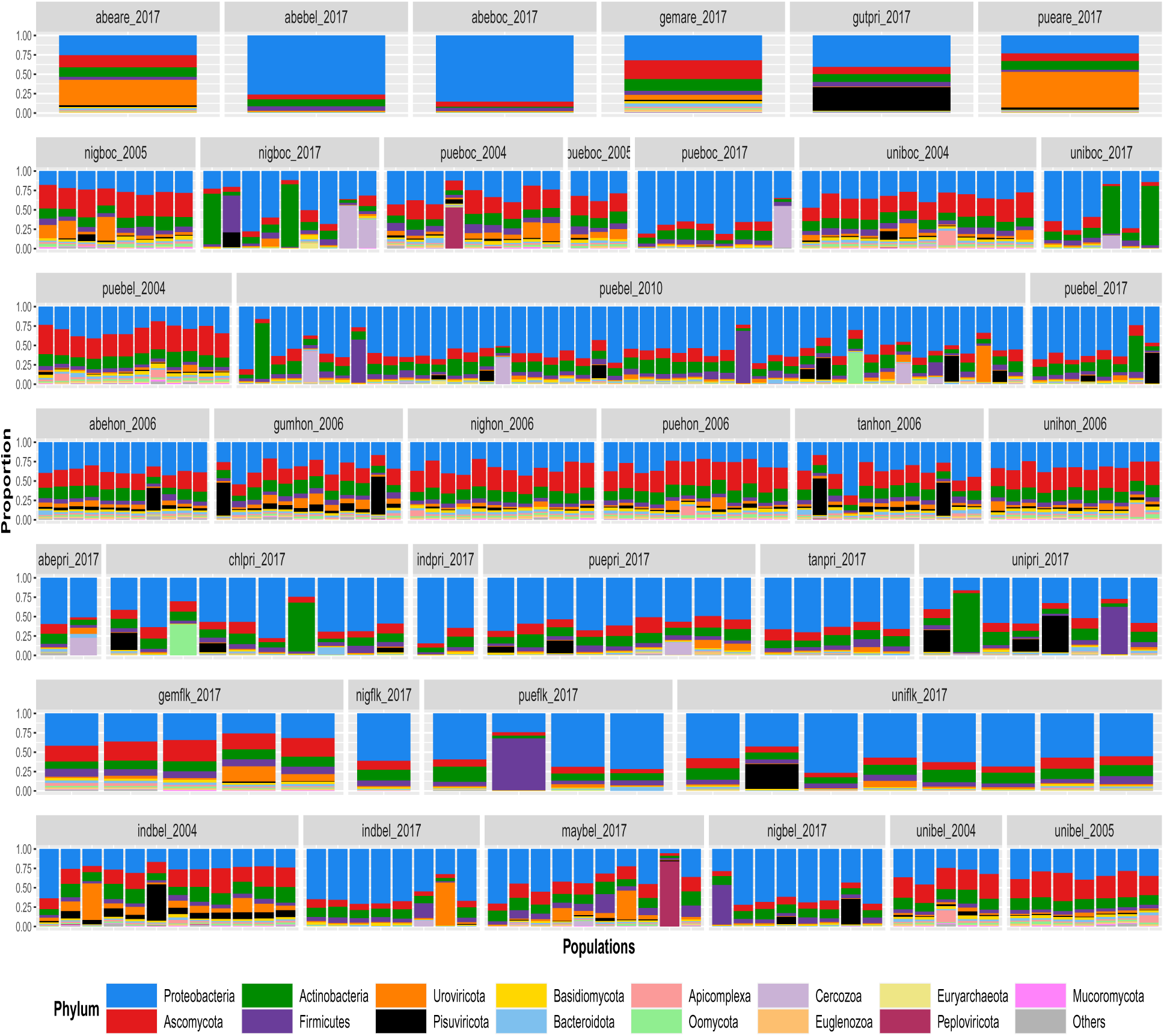
Taxonomic composition of the gill microbiome at the phylum level. Each stacked bar corresponds to one of the 304 samples. Taxonomic assignments are based on genomic read classification. The first three letters of the grey labels correspond to the host species (pue: *H. puella*, uni: *H. unicolor*, nig: *H. nigricans*, abe: *H. aberrans*, ind: *H. indigo*, chl: *H. chlorurus*, gum: *H. gummigutta*, gut: *H. guttavarius*, may: *H. maya*, gem: *H. gemma*, tan: *H. randallorum*, sp1: *H. sp.1*) and the last three letters to the location (are: Cayo Arenas, flk: Florida Keys, bel: Belize, hon: Honduras, boc: Bocas del Toro, pri: Puerto Rico).

**Suppl. Figure 2.**
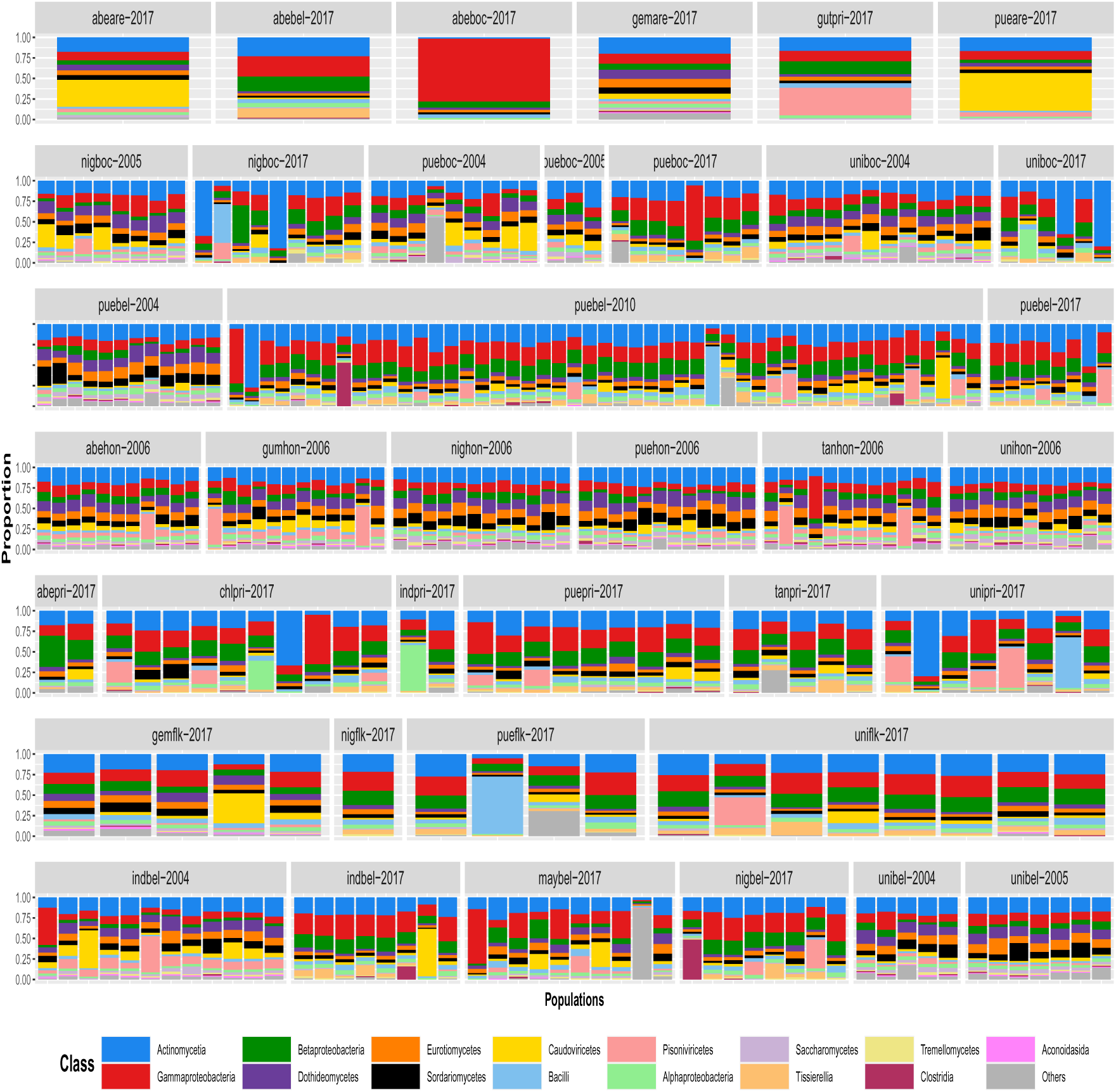
Taxonomic composition of the gill microbiome from read recruitment analysis at the class level. Each stacked bar corresponds to one of the 304 samples. Taxonomic assignments are based on genomic read classification. The first three letters of the grey labels correspond to the host species (pue: *H. puella*, uni: *H. unicolor*, nig: *H. nigricans*, abe: *H. aberrans*, ind: *H. indigo*, chl: *H. chlorurus*, gum: *H. gummigutta*, gut: *H. guttavarius*, may: *H. maya*, gem: *H. gemma*, tan: *H. randallorum*, sp1: *H. sp.1*) and the last three letters to the location (are: Cayo Arenas, flk: Florida Keys, bel: Belize, hon: Honduras, boc: Bocas del Toro, pri: Puerto Rico).

**Suppl. Figure 3.**
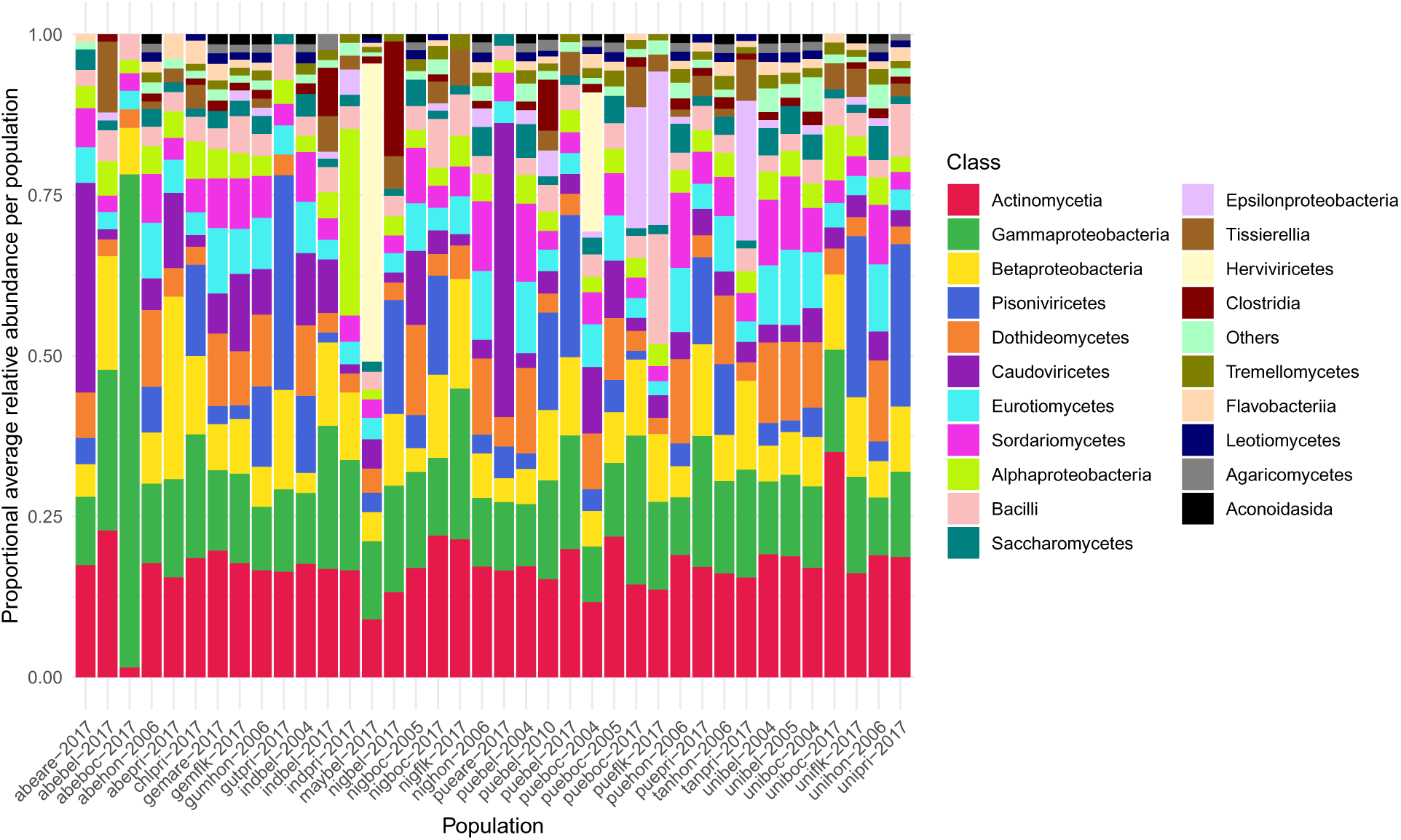
Proportional average relative abundance per population at the class level. The x-axis represents the population names (species/location). Proportions below a certain threshold (0.01 in this case) are filtered out.

**Suppl. Figure 4.**
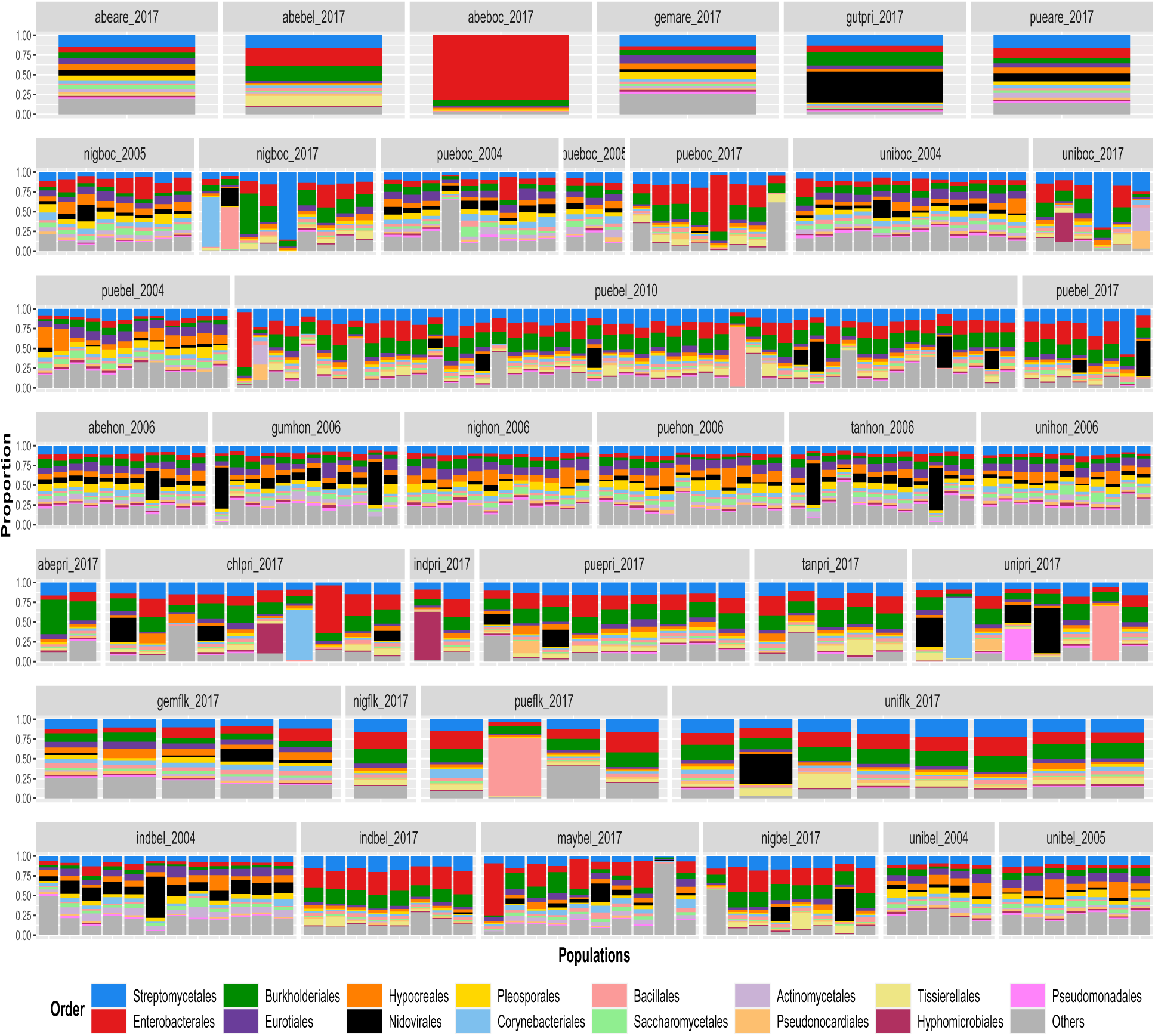
Taxonomic composition of the gill microbiome at the order level. Each stacked bar corresponds to one of the 304 samples. Taxonomic assignments are based on genomic read classification. The first three letters of the grey labels correspond to the host species (pue: *H. puella*, uni: *H. unicolor*, nig: *H. nigricans*, abe: *H. aberrans*, ind: *H. indigo*, chl: *H. chlorurus*, gum: *H. gummigutta*, gut: *H. guttavarius*, may: *H. maya*, gem: *H. gemma*, tan: *H. randallorum*, sp1: *H. sp.1*) and the last three letters to the location (are: Cayo Arenas, flk: Florida Keys, bel: Belize, hon: Honduras, boc: Bocas del Toro, pri: Puerto Rico).

**Suppl. Figure 5.**
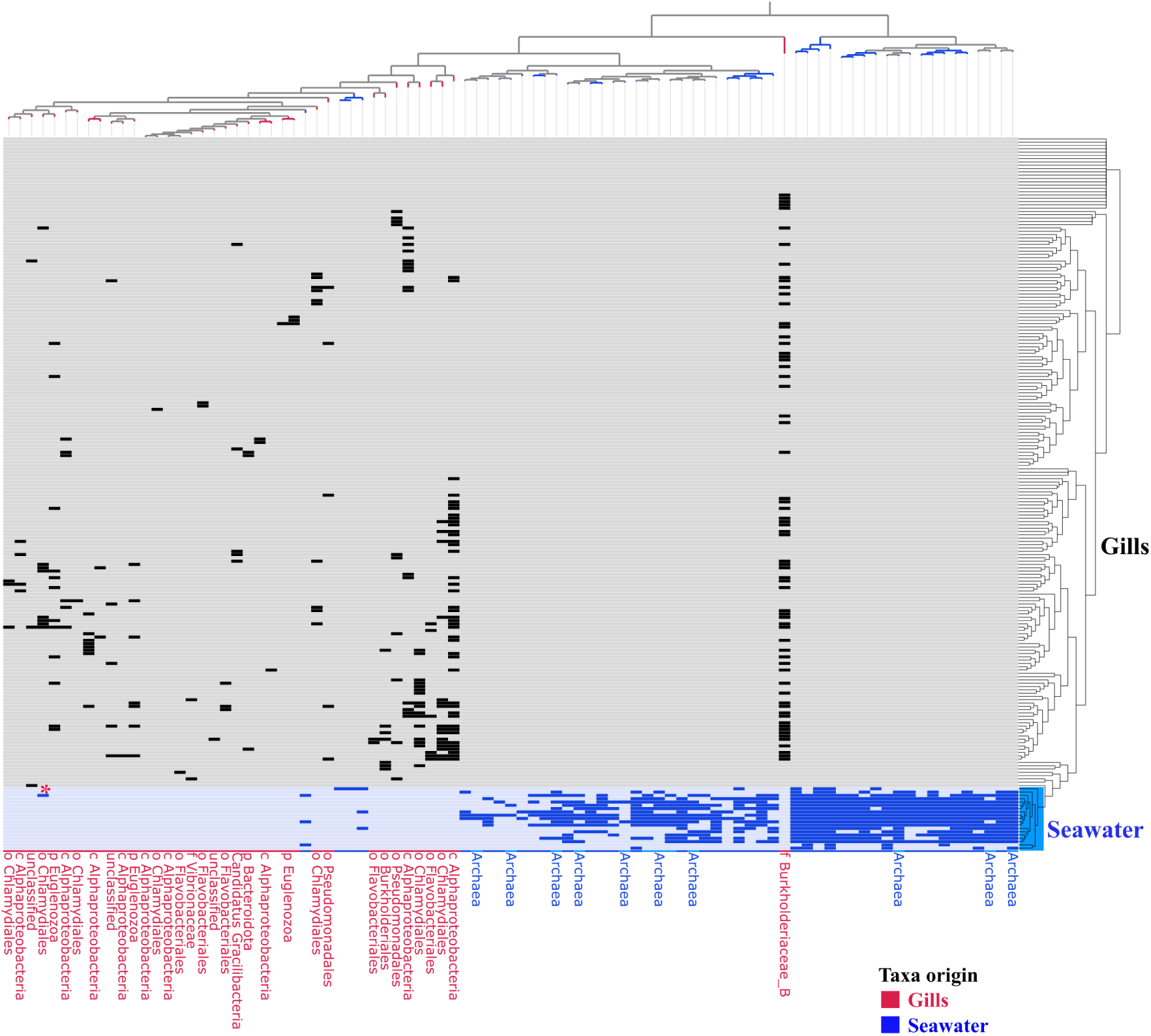
Presence/absence heatmap of *rpL13* across gill and seawater samples. The rows correspond to metagenomic samples, columns are *rpL13* representative sequences from 95% nucleotide clustering, and the data points in the heatmap represent presence/absence of *rpL13* in the metagenomes. The metagenomes (rows) are organized by hierarchical clustering based on the detection of *rpL13* while the columns are organized by *rpL13* co-occurence across samples. *rpL13* was used as a proxy for the detection microbial populations in metagenomes. Gill derived *rpL13* were only detected in gill metagenomes and not seawater samples and vice versa, highlighting that the gills have a distinct microbiome. Red star denotes Chlamydiales population found in both gills and water column. Each leaf of the tree is decorated with metadata denoting origin of microbial population: (red: gill taxa; blue: water column taxa). We used an anvi’o detection threshold of 0.7 to define the presence of a ribosomal protein in a metagenome with read recruitment.

**Suppl. Figure 6.**
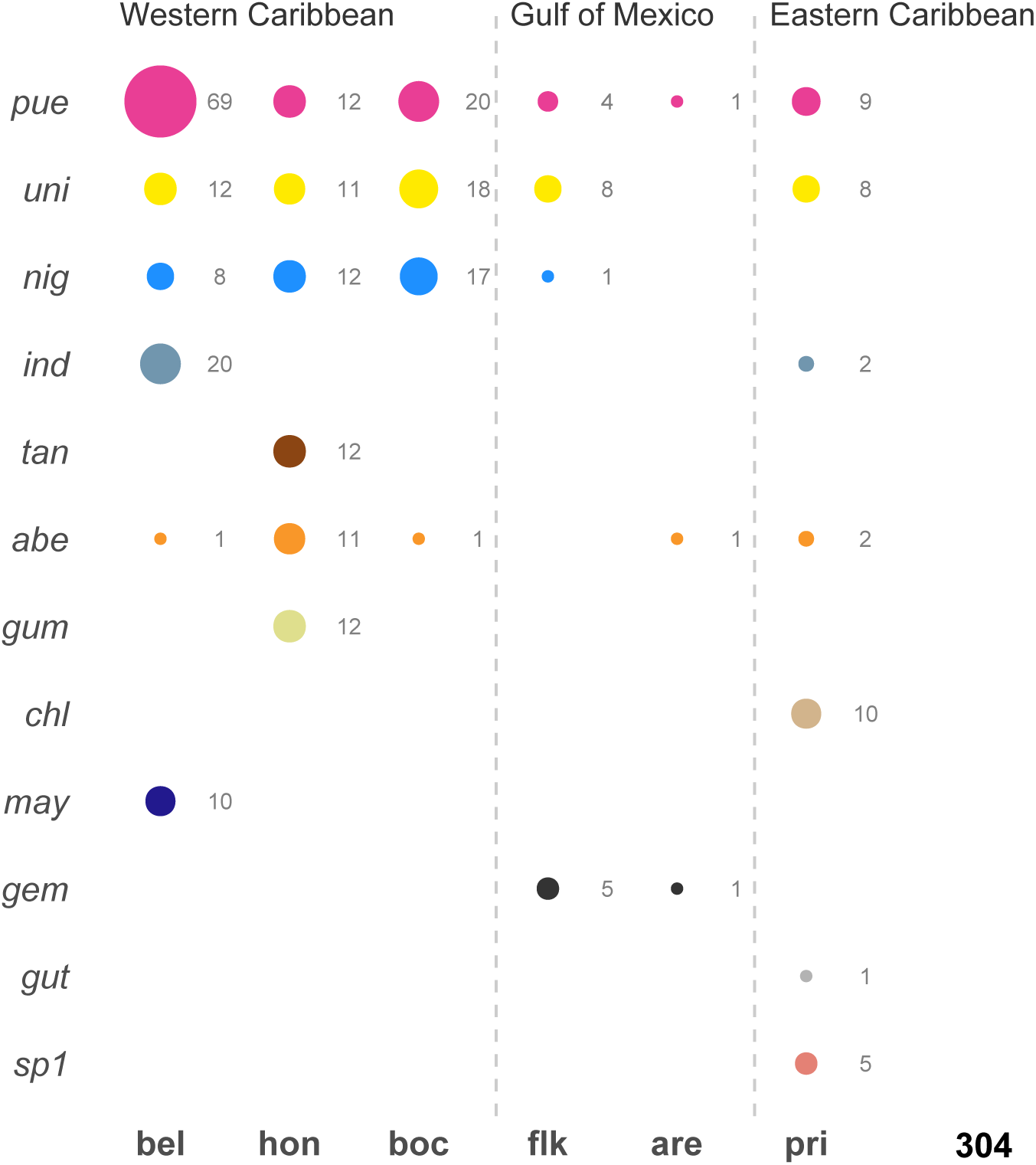
**Overview of the sampling design**. It includes 304 shotgun-sequenced gill samples from 12 hamlet species – pue: *Hypoplectrus puella*, uni: *Hypoplectrus unicolor*, nig: *Hypoplectrus nigricans*, abe: *Hypoplectrus aberrans*, ind: *Hypoplectrus indigo*, chl: *Hypoplectrus chlorurus*, gum: *Hypoplectrus gummigutta*, gut: *Hypoplectrus guttavarius*, may: *Hypoplectrus maya*, gem: *Hypoplectrus gemma*, tan: *Hypoplectrus randallorum*, sp1: *Hypoplectrus sp.1* were collected at 6 locations [Cayo Arenas (are) and the Florida Keys (flk) in the Gulf of Mexico, Belize (bel), Honduras (hon), Bocas del Toro in Panama (boc), and Puerto Rico (pri)].

**Suppl. Figure 7.**
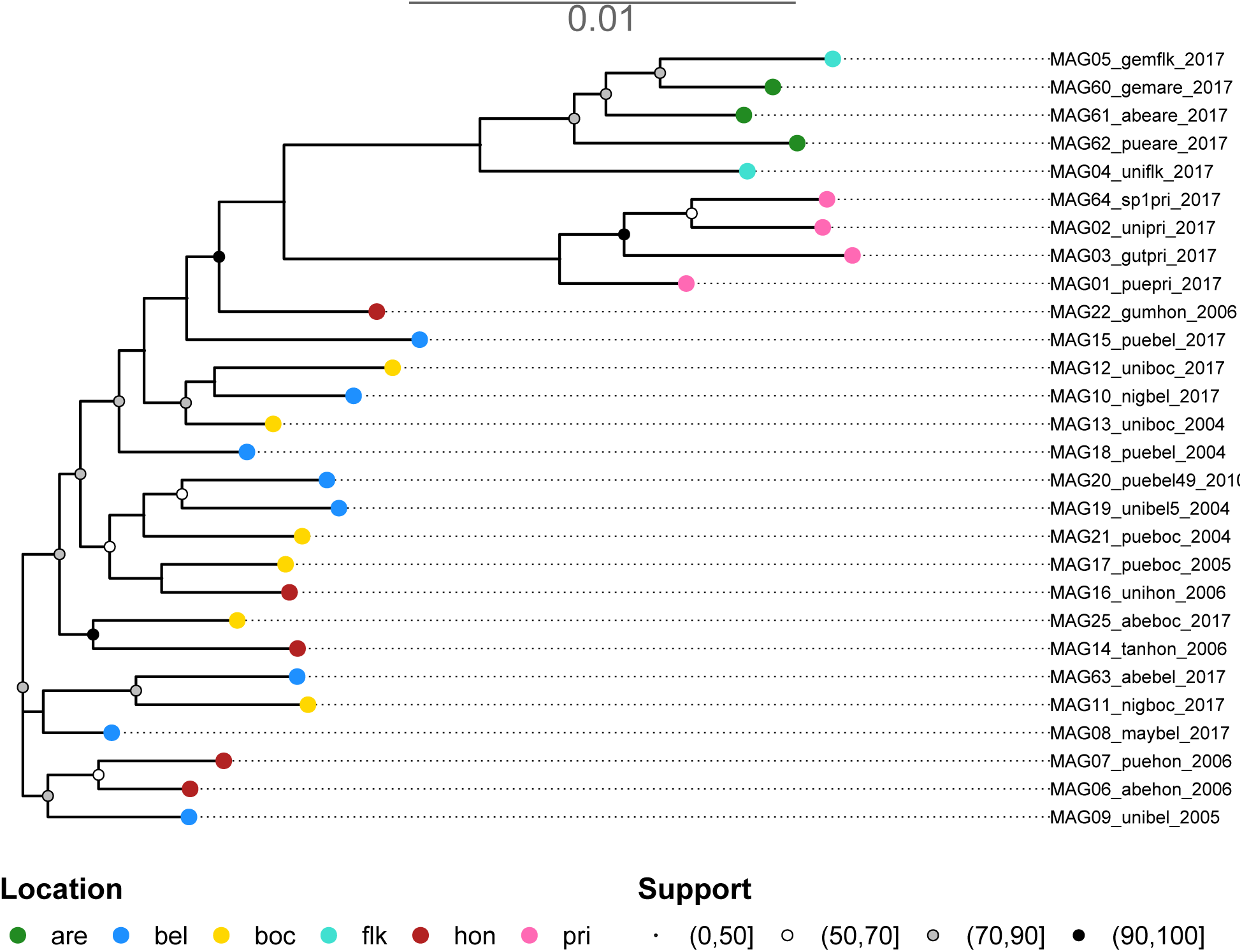
Phylogenetic tree of 28 MAGs from the most prevalent taxon in the Burkholderiaceae-B family. The MAGs from Cayo Arenas (are) and the Florida Keys (flk) in the Gulf of Mexico, and Puerto Rico (pri) formed a clade diverged from the remaining locations: Belize (bel), Honduras (hon) and Bocas del Toro in Panama (boc).

**Suppl. Figure 8.**
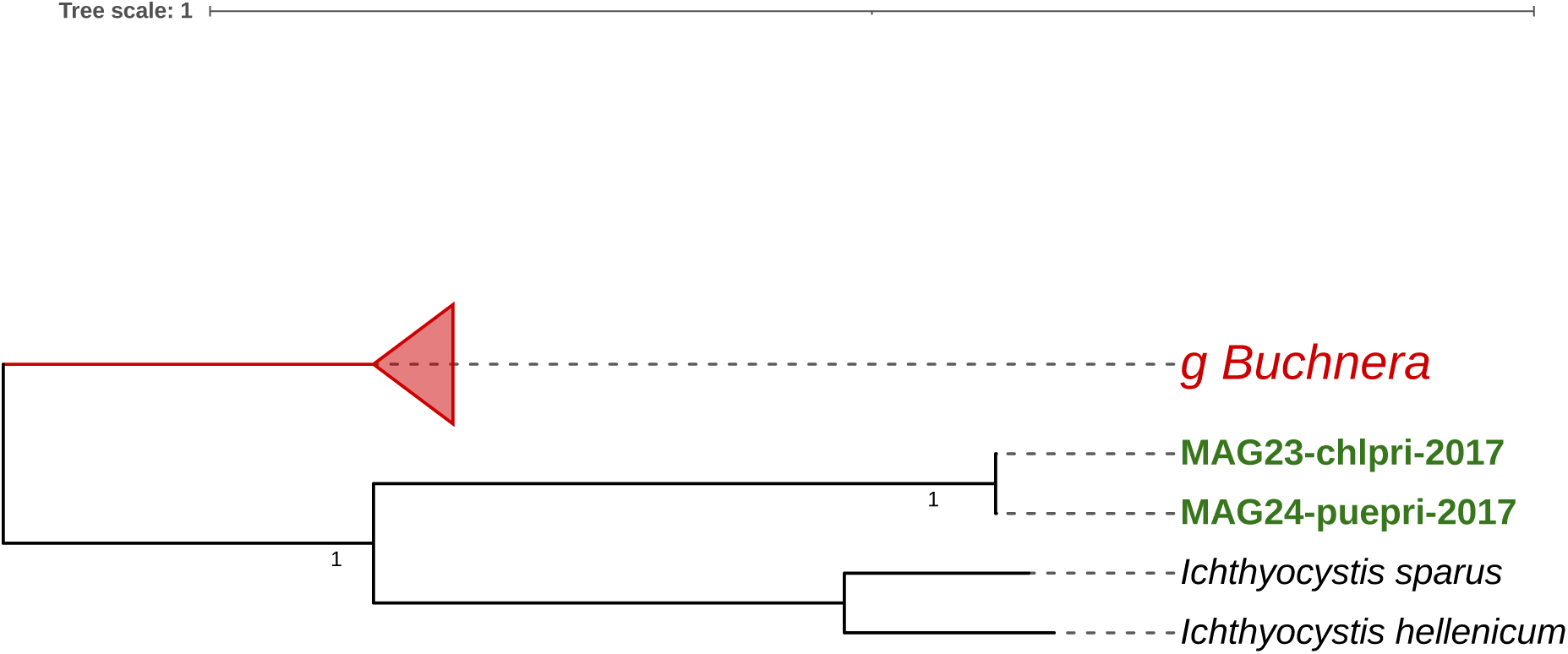
Phylogentic placement of the MAGs in the 2013Ark19i family (order Burkholderiales). The tree is rooted with the genus *Buchnera* (in red). Support values at the nodes represent non-parametric bootstrap values.

**Suppl. Figure 9.**
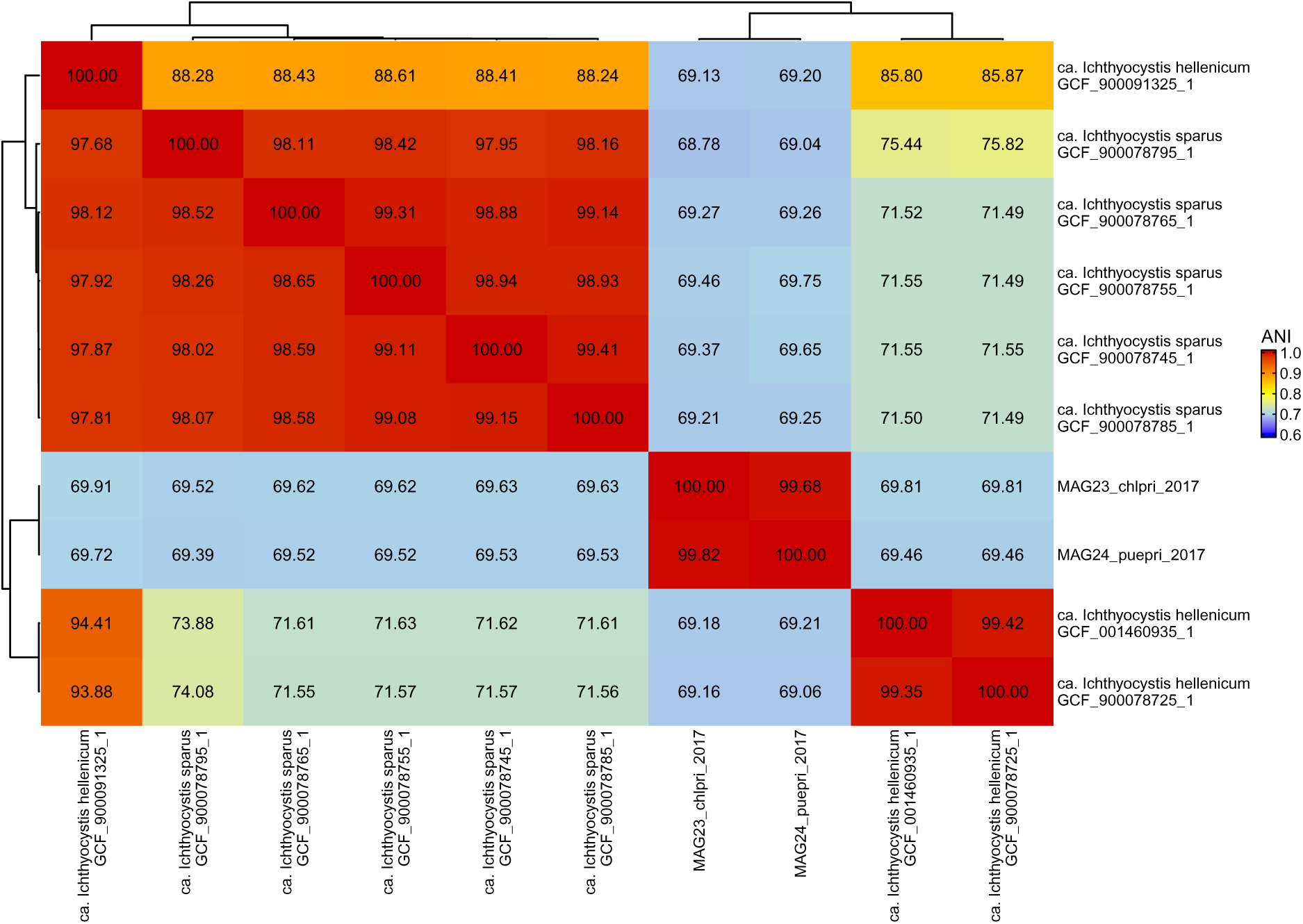
Average Nucleotide Identity (ANI) of the MAGs in the 2013Ark19i family (order Burkholderiales). These two MAGs were very closely related (>99% ANI) but showed an ANI <70% with the other genomes in the family (*Ichthyocystis sparus* and *hellenicum*), which are known marine fish gill pathogens. The analysis included the two MAGs recovered in this family and the closest representative relatives retrieved from NCBI. The genomes are grouped based on hierarchical clustering applied to the ANI values.

**Suppl. Figure 10.**
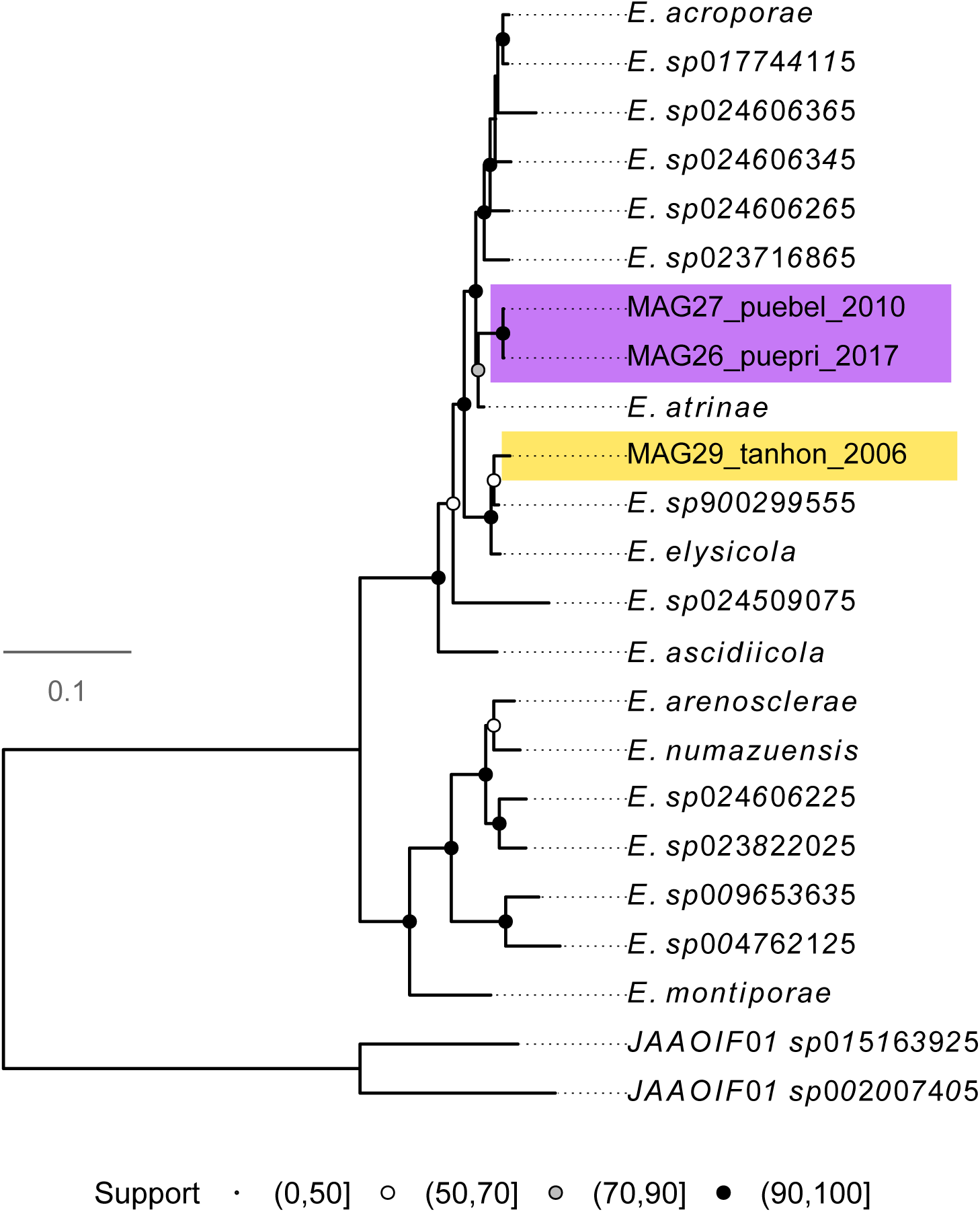
Phylogenomic placement of the MAGs in the family Endozoicomonadaceae (order Pseudomonadales). The three MAGs clustered with the genus *Endozoicomonas* – two with *E. atrinae*, which was isolated from the gut of the comb pen shell (*Atrina pectinata*), and one with *E. sp900299555* and *E. elysicola*, two fish gill and skin pathogens. Support symbols at the nodes represent non-parametric bootstrap values.

**Suppl. Figure 11.**
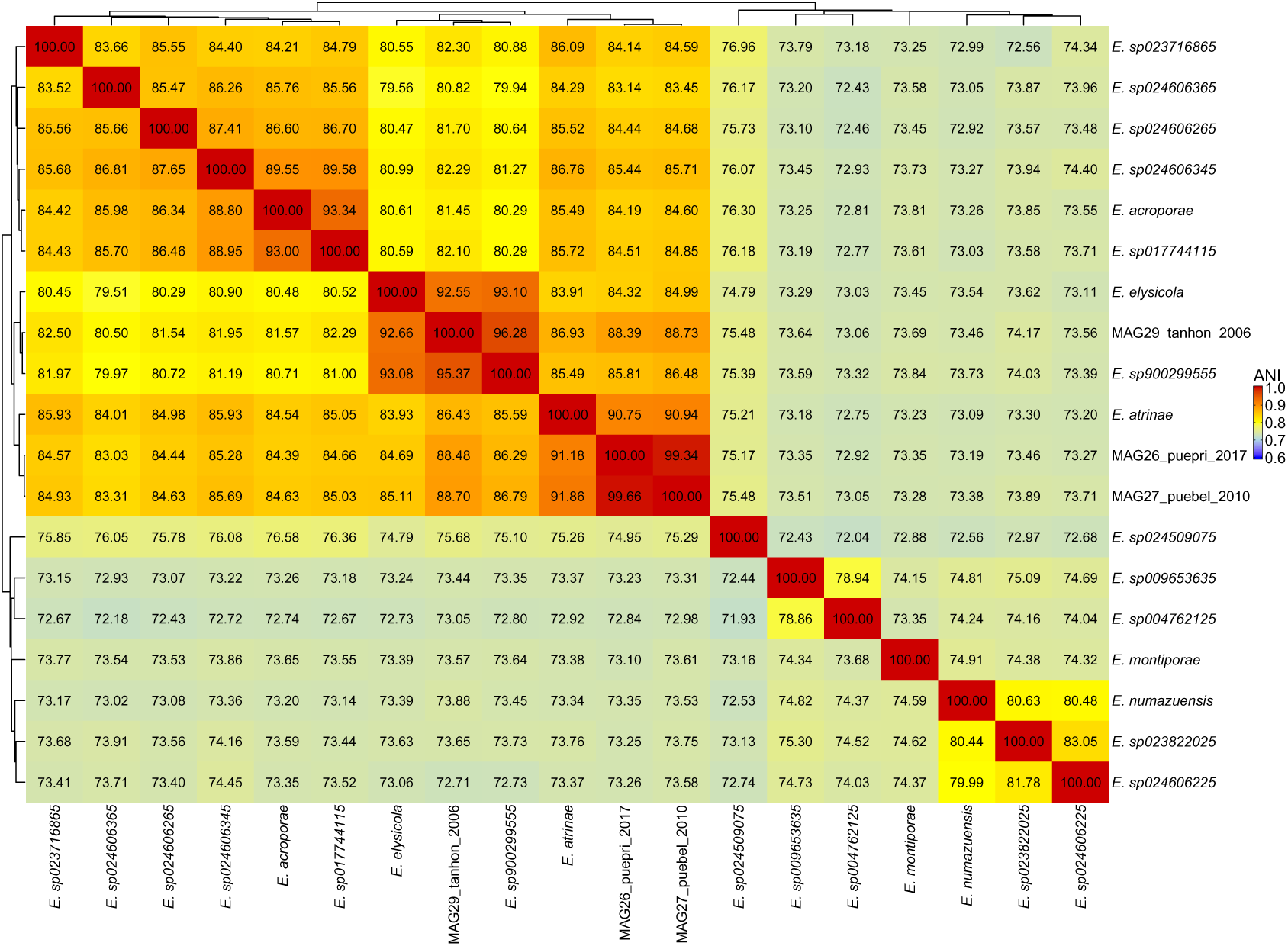
Average Nucleotide Identity (ANI) of the MAGs in the family Endozoicomonadaceae (order Pseudomonadales). Two MAGs had a highest ANI of 92% with *Endozoicomonas atrinae*, which was isolated from the gut of the comb pen shell (*Atrina pectinata*), and one had a higherst ANI of >96% with *E. sp900299555* and *E. elysicola*, two marine fish gill and skin pathogens. The analysis included the three MAGs recovered in this family and the closest representative relatives retrieved from NCBI. The genomes were grouped based on hierarchical clustering applied to the ANI values.

**Suppl. Figure 12.**
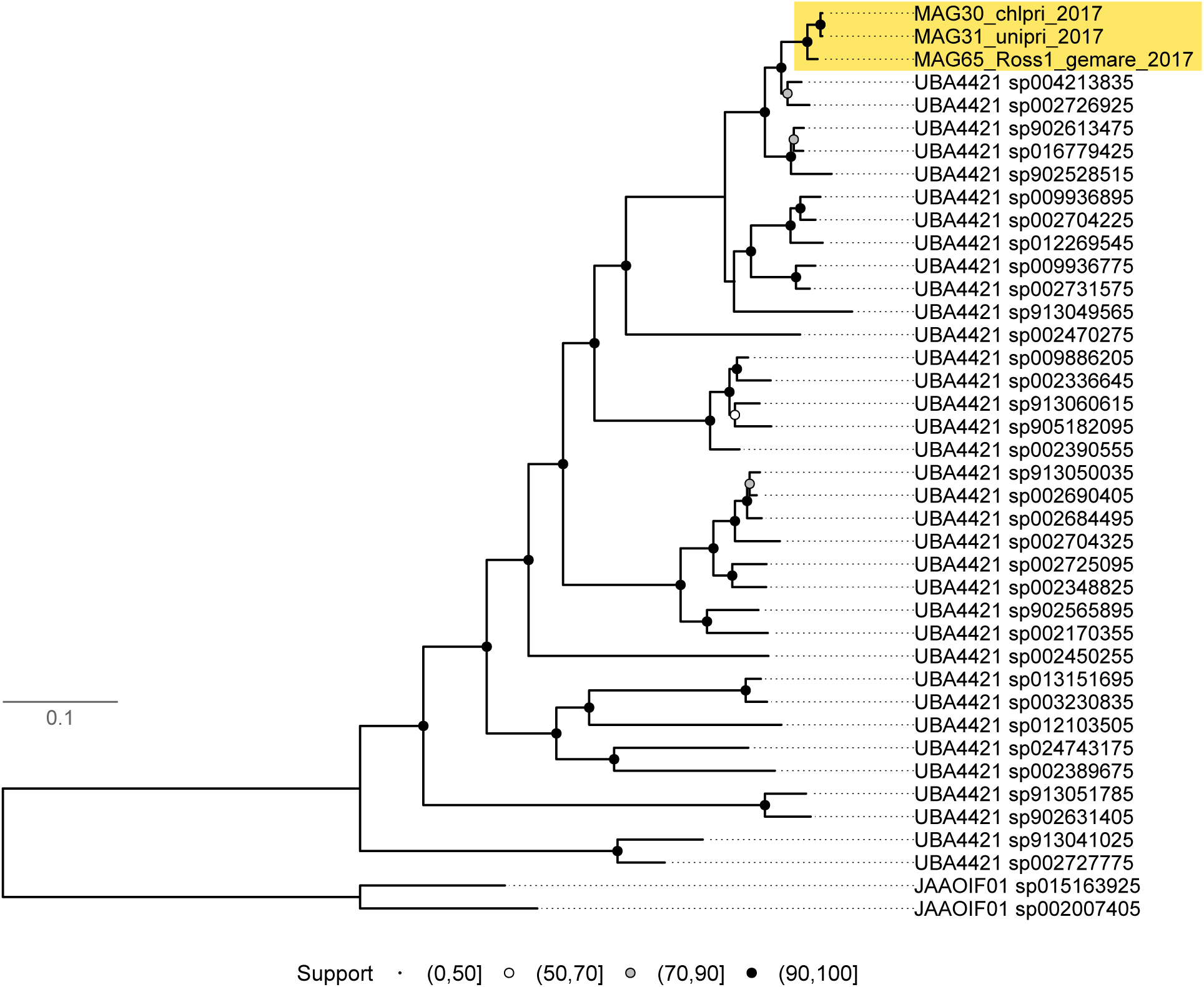
Phylogenomic placement of the MAGs in the family HTCC2089 (order Pseudomonadales). These MAGs clustered with species in the genus *UBA4421*, in particular with *UBA4421 sp004213835* and *UBA4421 sp002726925* that were isolated from seawater in the Mediterranean Sea and Pacific Ocean, respectively. Support symbols at the nodes represent non-parametric bootstrap values.

**Suppl. Figure 13.**
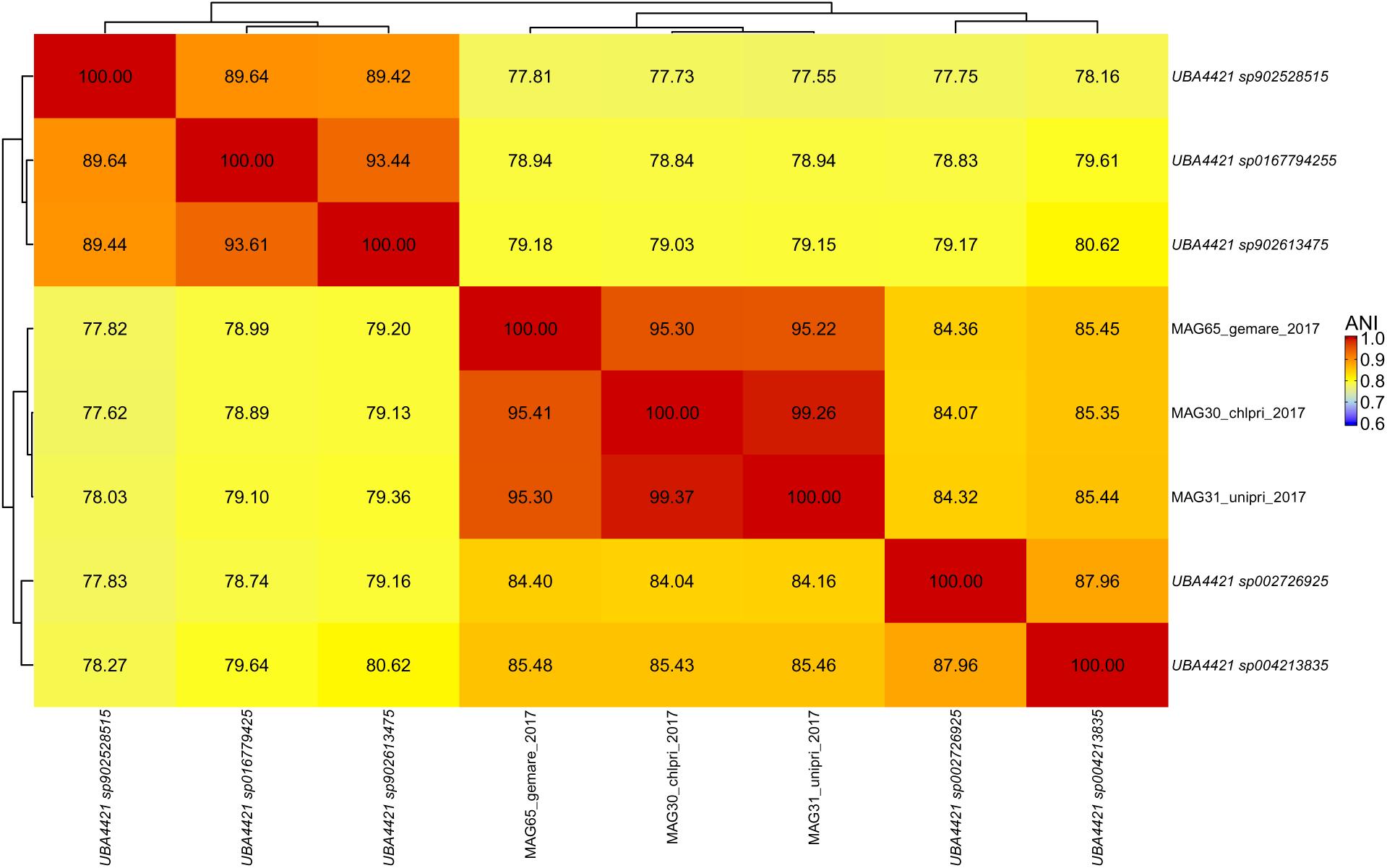
**Average Nucleotide Identity (ANI) of the MAGs in the family HTCC2089 (order Pseudomonadales)**. The average ANI between the MAGs and their closest relatives, *UBA4421 sp004213835* and *UBA4421 sp002726925*, was 85%. These relatives were isolated from seawater in the Mediterranean Sea and Pacific Ocean, respectively. The analysis included the three MAGs recovered in this family and the closest representative relatives retrieved from NCBI. The genomes were grouped based on hierarchical clustering applied to the ANI values.

**Suppl. Figure 14.**
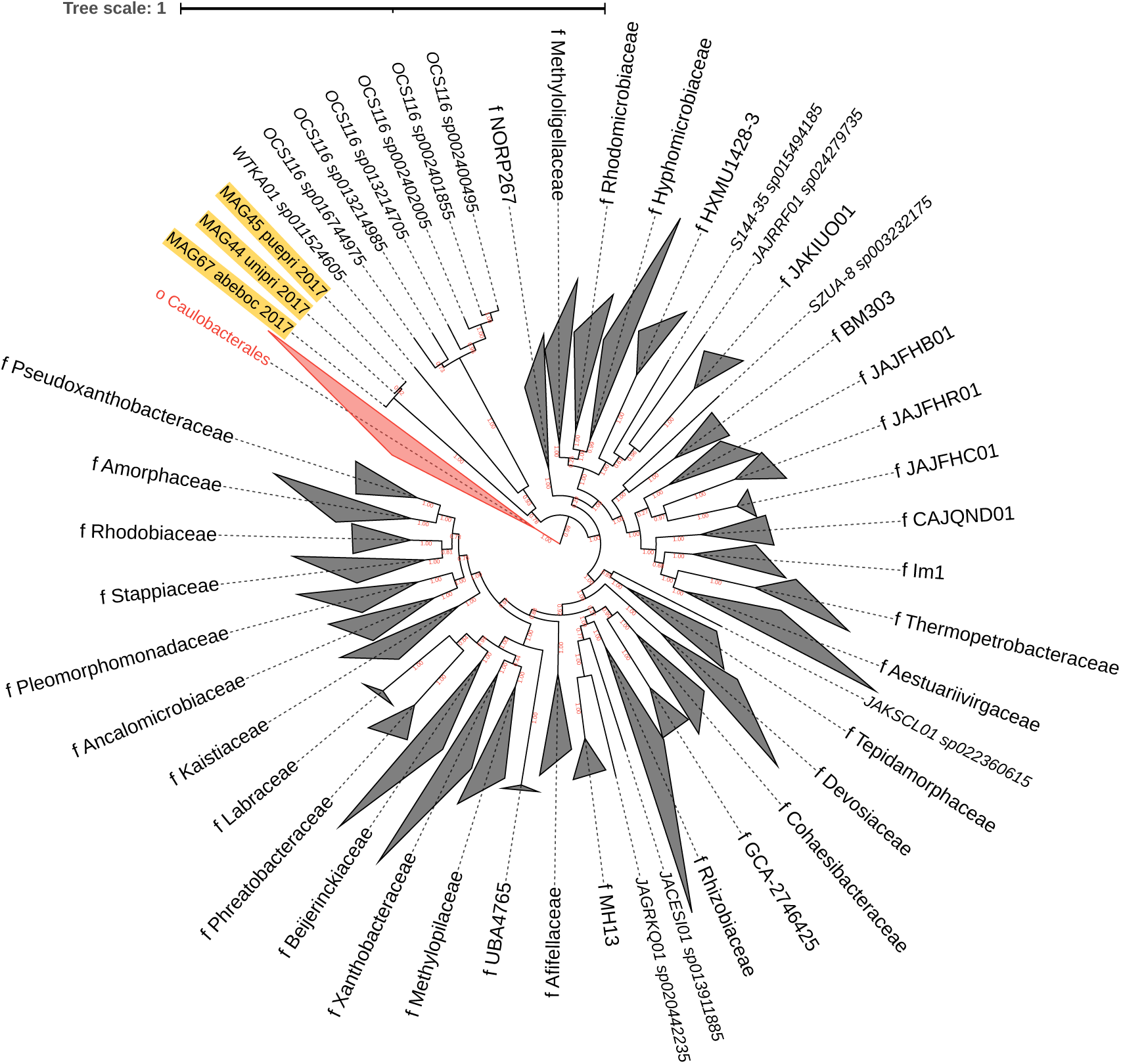
Phylogenomic placement of the three MAGs in the order Rhizobiales. These MAGs (highlighted) form a distinct clade that is sister to the WTKA01 family. Family names are indicated by a leading "f" where family nodes were collapsed. Triangles size represent the number of genomes in these collapsed nodes. The tree is rooted with the order *Caulobacterales* (in red). Support values at the nodes represent non-parametric bootstrap values.

**Suppl. Figure 15.**
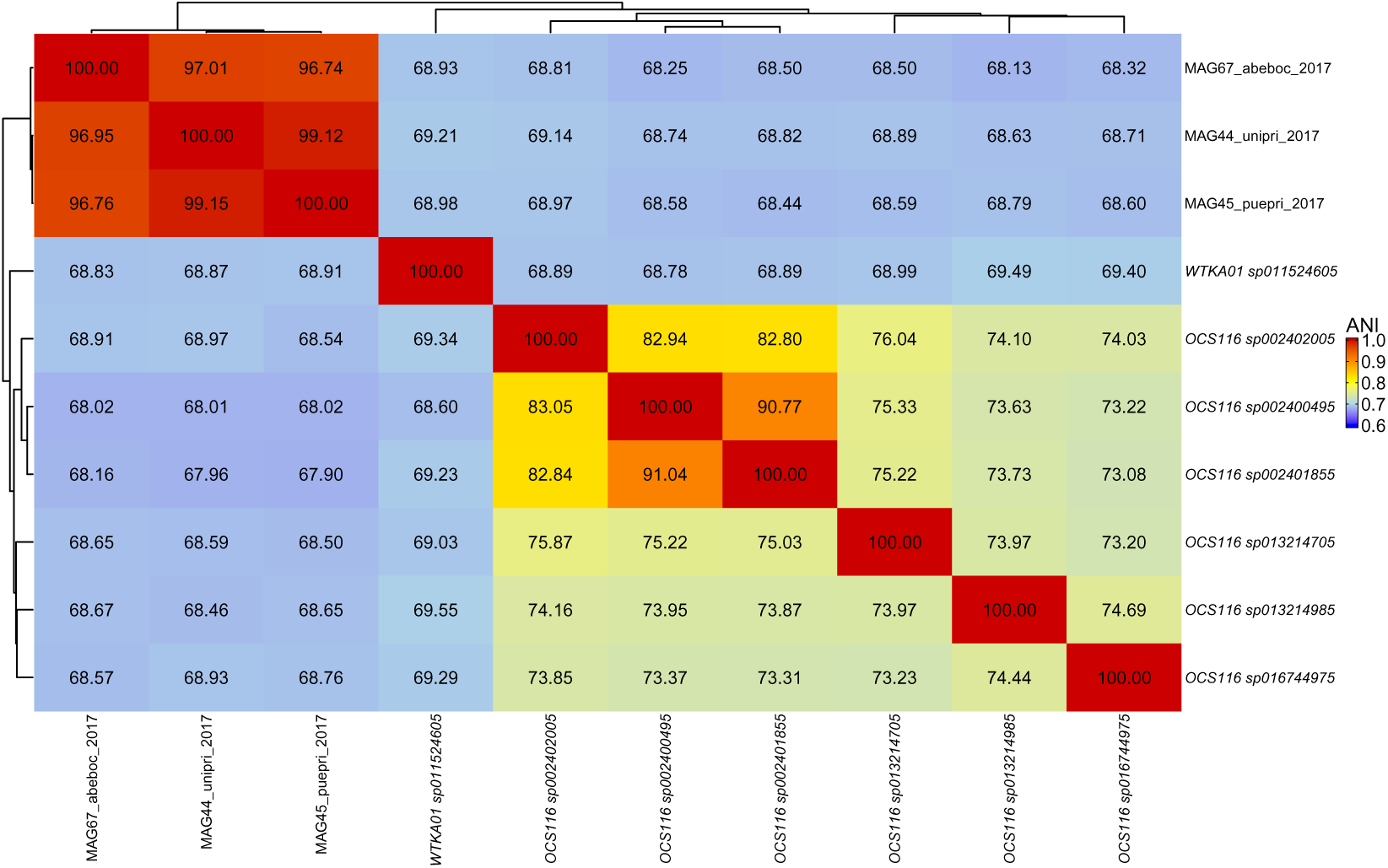
Average Nucleotide Identity (ANI) of the three MAGs in the order Rhizobiales. The MAGs had highest ANI of 69% with *WTKA01 sp011524605* that were isolated from coral reef macroalgae biofilm. The analysis included the three MAGs recovered in this order and the closest representative relatives retrieved from NCBI. The genomes were grouped based on hierarchical clustering applied to the ANI values.

**Suppl. Figure 16.**
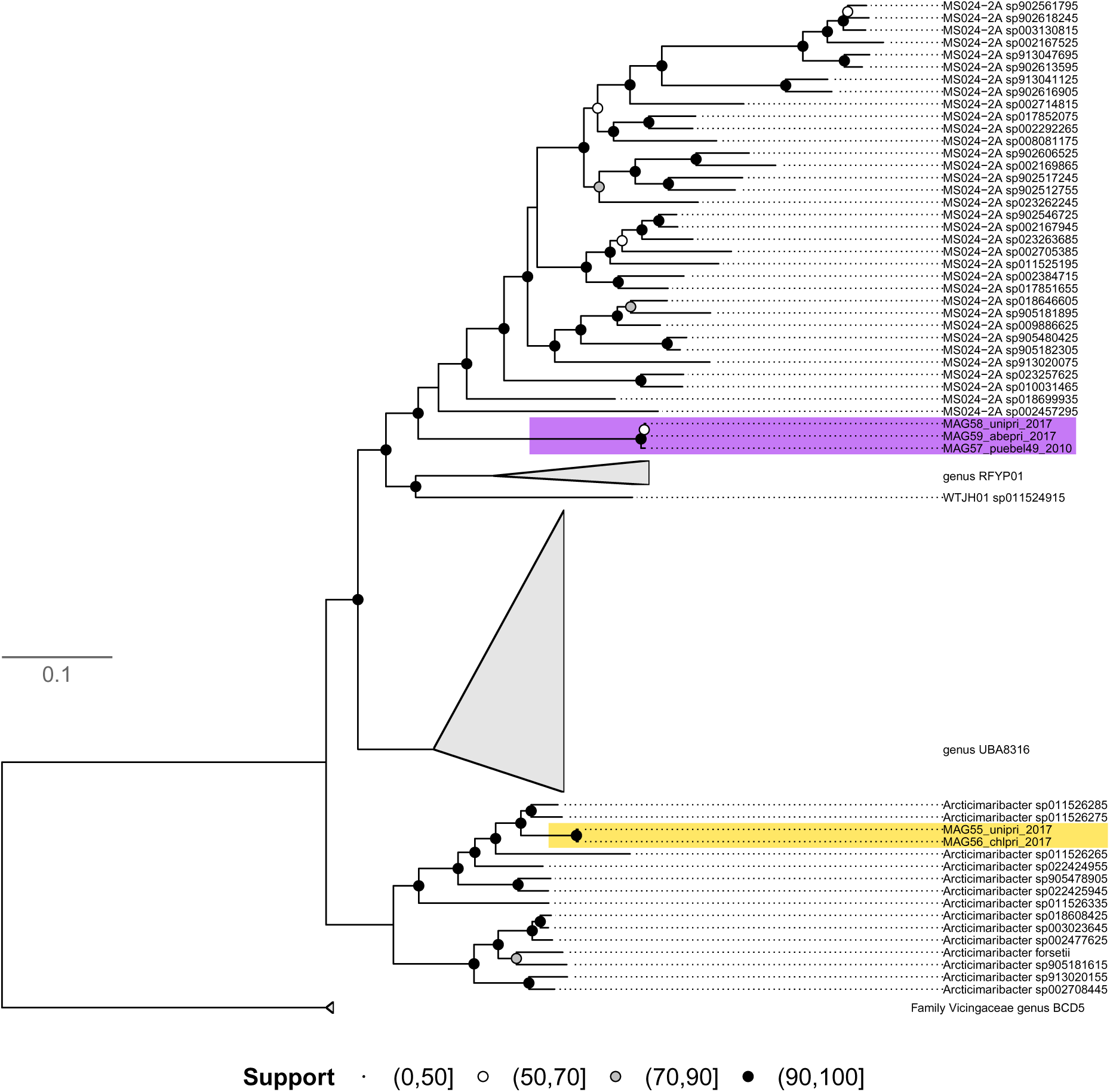
Phylogenomic placement of the five MAGS in the order Flavobacteriales. These MAGS belong to the family Flavobacteriaceae, but to two different clades (general genus *MS024-2A* and *Arcticimaribacter*). These two groups are mainly represented by seawater taxa. Support symbols at the nodes represent non-parametric bootstrap values

**Suppl. Figure 17.**
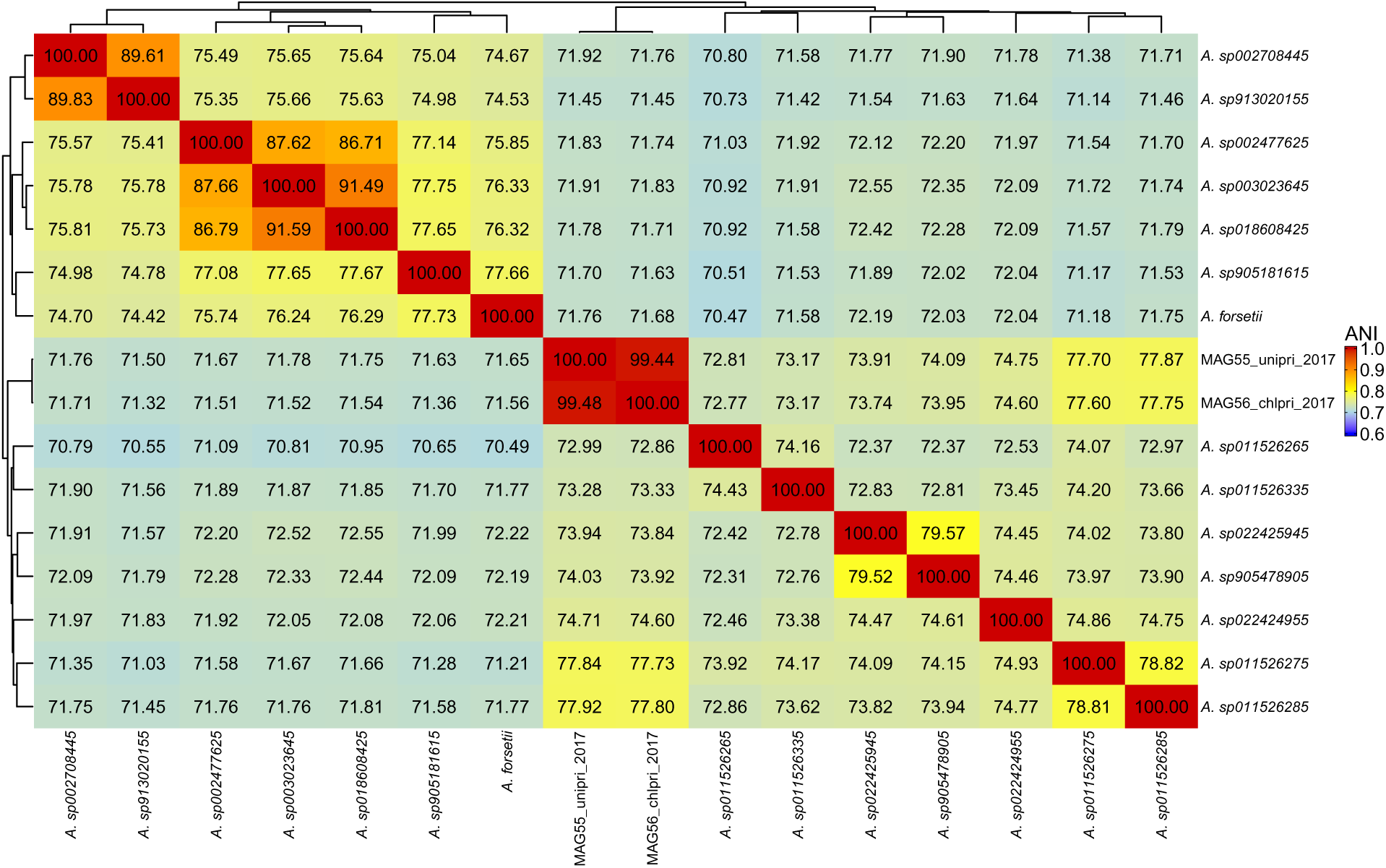
Average Nucleotide Identity (ANI) of the two MAGs in the family Flavobacteriaceae, genus *Arcticimaribacter*. *Arcticimaribacter* is abbreviated as *A.* The MAGs had highest ANI of 78% with *A. sp011526285* that was isolated from coral reef seawater. The analysis included the two MAGs recovered in this genus and the closest representative relatives retrieved from NCBI. The genomes were grouped based on hierarchical clustering applied to the ANI values.

**Suppl. Figure 18.**
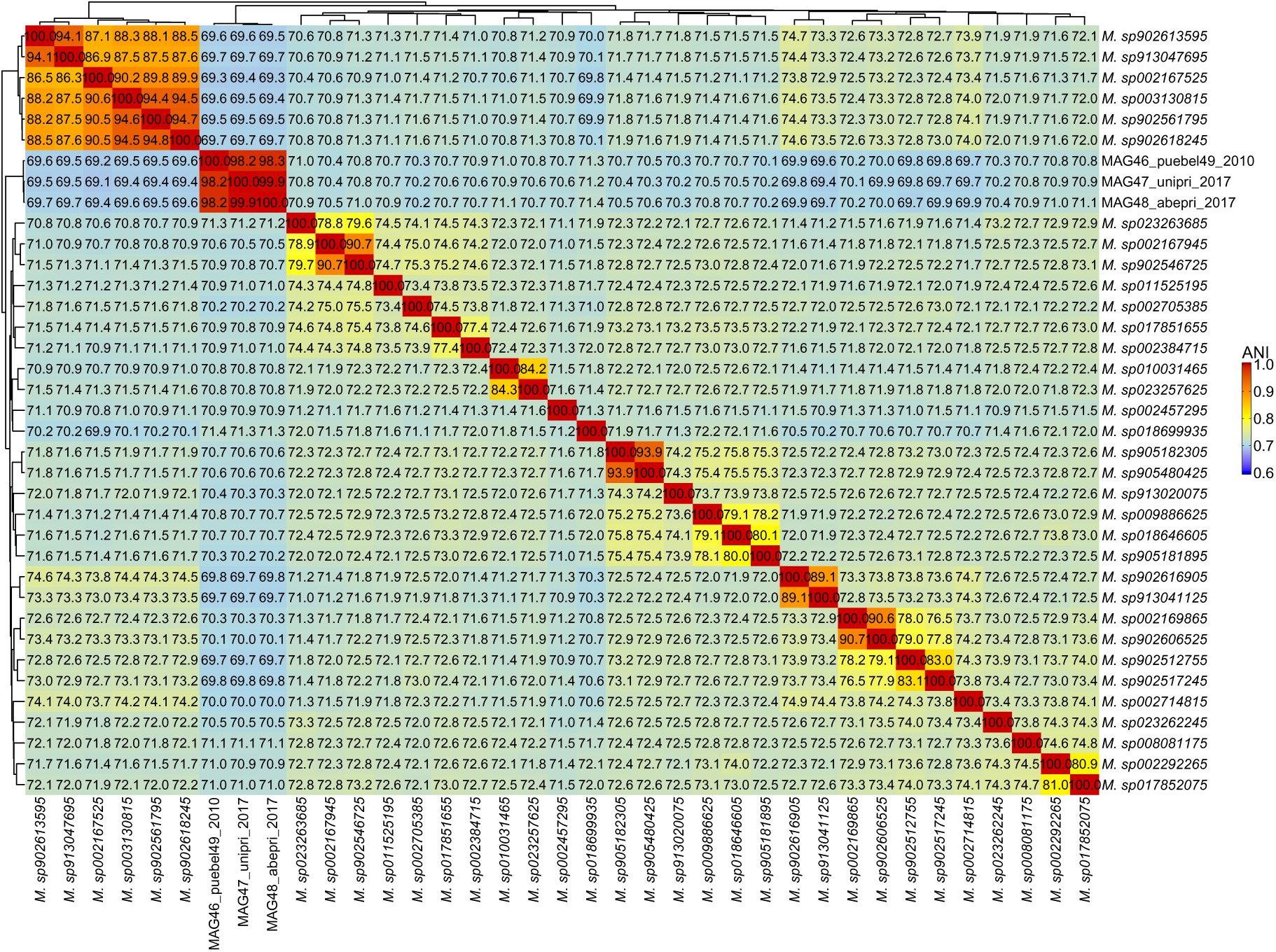
Average Nucleotide Identity (ANI) of the three MAGs in the family Flavobacteriaceae, genus *MS024-2A*. *MS024-2A* is abbreviated as *M.* The MAGs had an average ANI of 71.3% with *M. sp018699935* that was isolated from hypoxic seawater. The analysis included the three MAGs recovered in this genus and the closest representative relatives retrieved from NCBI. The genomes were grouped based on hierarchical clustering applied to the ANI values.

**Suppl. Figure 19.**
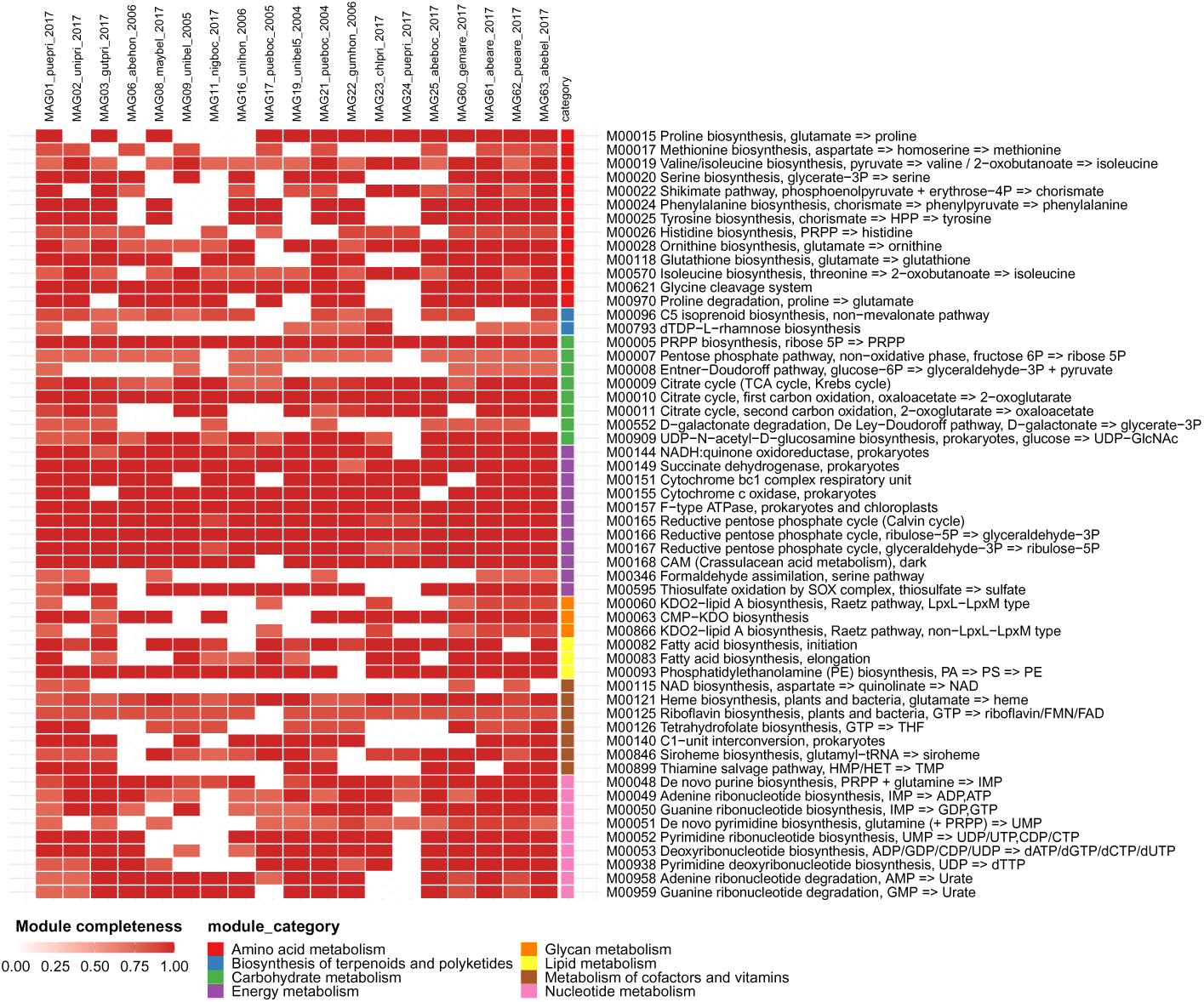
**Completeness of KEGG metabolic modules in the order Burkholderiales**. All MAGs belong to the most prevalent taxon in the Burkholderiaceae-B family except for MAG23-MAG24 relate to genus *Ichthyocystis* fish gill pathogens in family *2013Ark19i*. The color scale represents module completeness, from 0 to 1. Only modules with completeness ≥ 75% in at least one MAG are presented

**Suppl. Figure 20.**
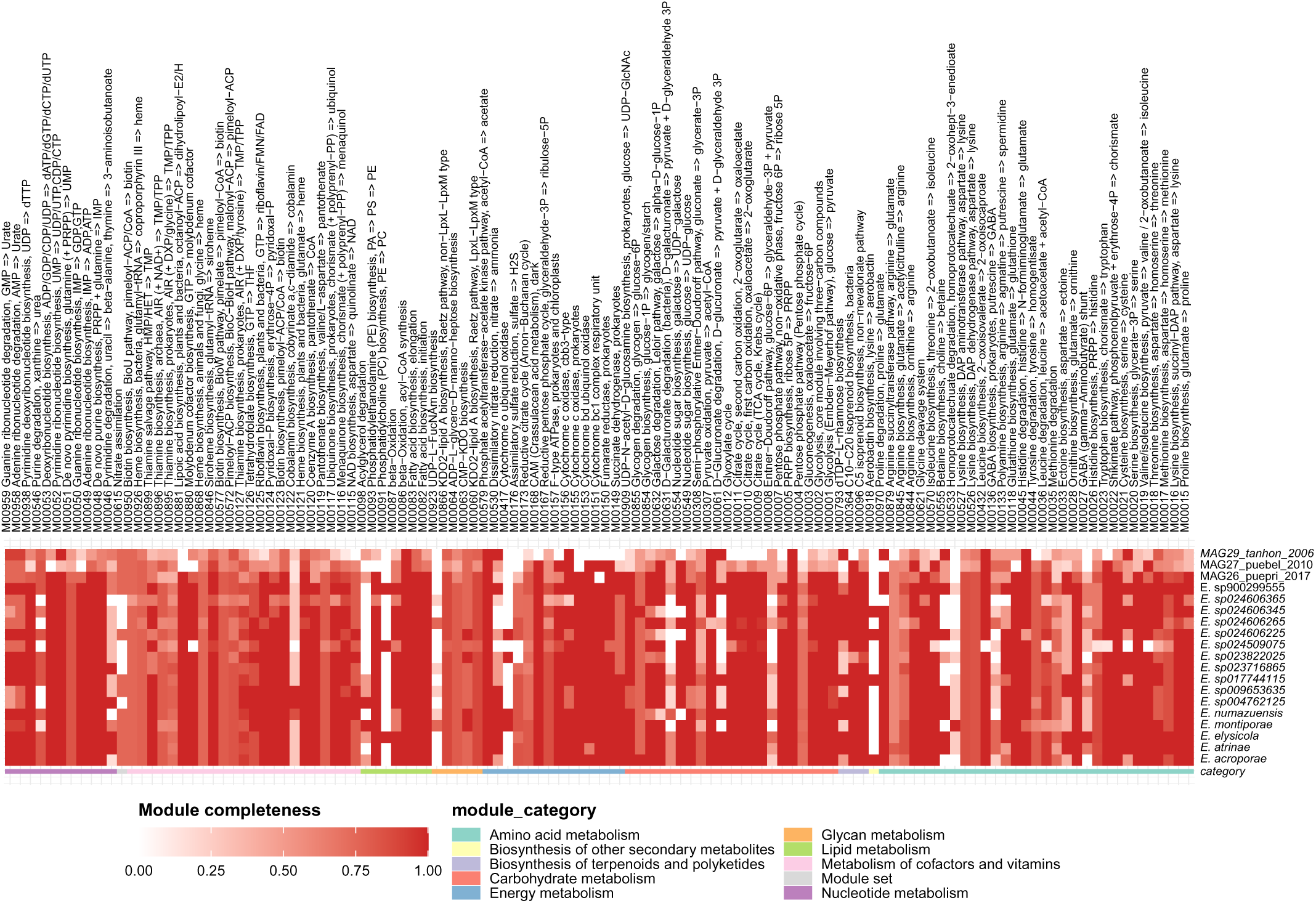
Completeness of KEGG metabolic modules in the genus Endozoicomonas (order Pseudomonadales). MAGs 26-27 relate to a species from the gut of the comb pen shell, and MAG29 to fish gill pathogens (but note that it has low completeness). The color scale represents module completeness, from 0 to 1. Only modules with completeness ≥ 75% in at least one genome are presented.

**Suppl. Figure 21.**
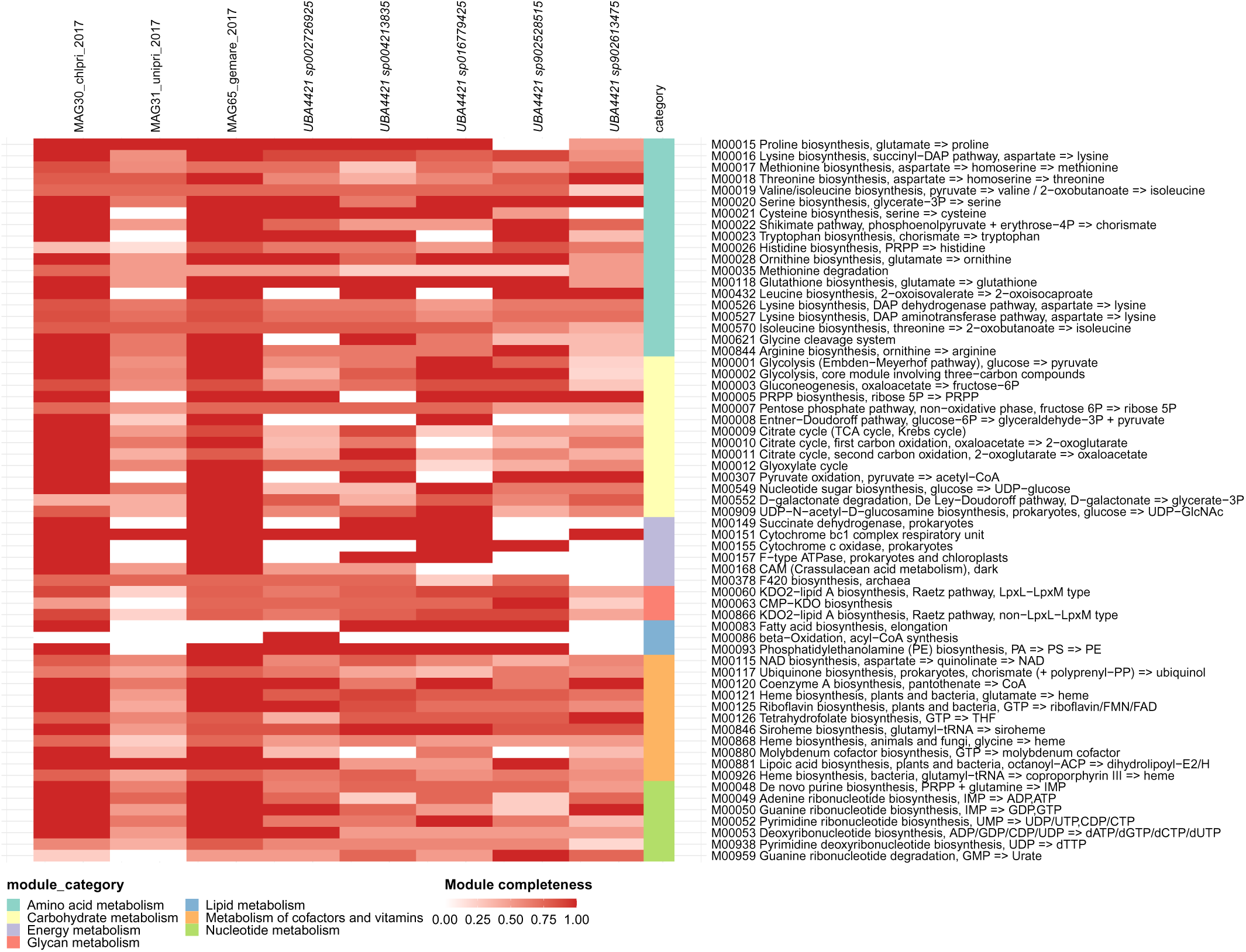
Completeness of KEGG metabolic modules in the genus *UBA4421* (order Pseudomonadales). Three MAGs (30, 31 and 65) were recovered in this genus, which relates to seawater taxa. The color scale represents module completeness, from 0 to 1. Only modules with completeness ≥ 75% in at least one genome are presented.

**Suppl. Figure 22.**
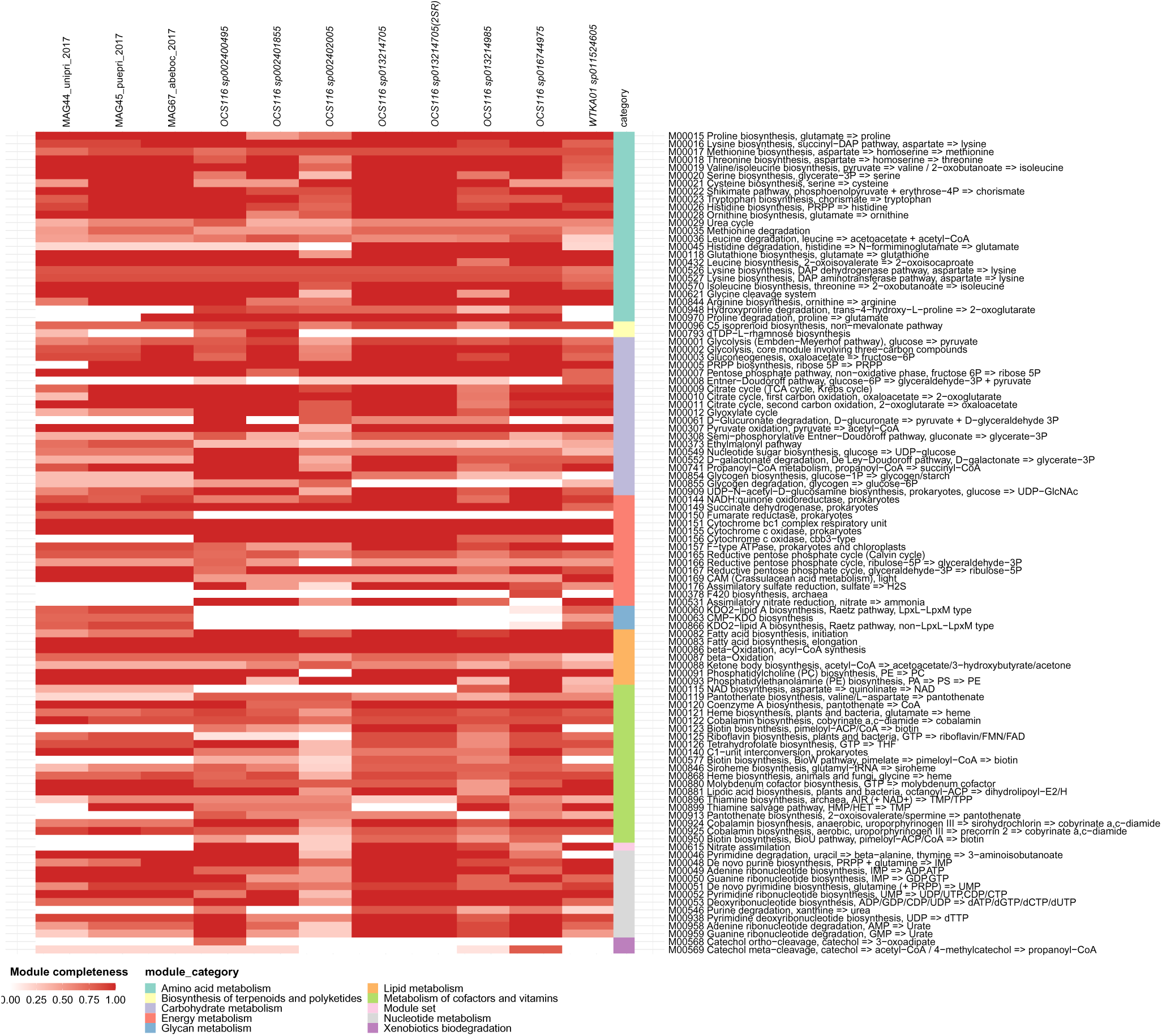
Completeness of KEGG metabolic modules in the genus *WTKA01* (order Rhizobiales). Three MAGs (44, 45, 67) were recovered in this genus, which relate to a species associated with coral reef macroalgae biofilm. 2SP means this genome is the second species representative (GCA_022968955_1) for the same species according to GTDB. The color scale represents module completeness from 0 to 1. Only modules with completeness ≥ 75% **^6^**i**^9^**n at least one genome are presented.

**Suppl. Figure 23.**
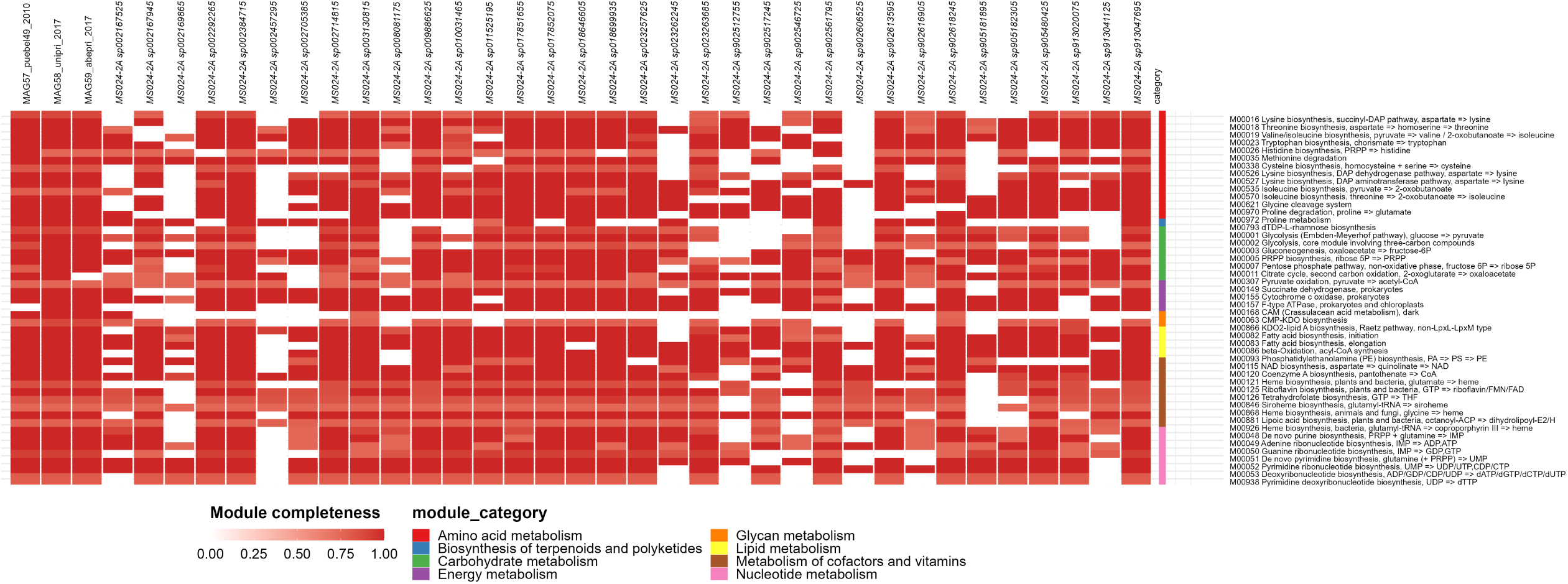
Completeness of KEGG metabolic modules in the genus *MS024-2A* (order Flavobacteriales). Three MAGs (57-59) were recovered in this genus, which relate to free-living seawater species. The color scale represents module completeness from 0 to 1. Only modules with completeness ≥ 75% in at least one genome are presented.

**Suppl. Figure 24.**
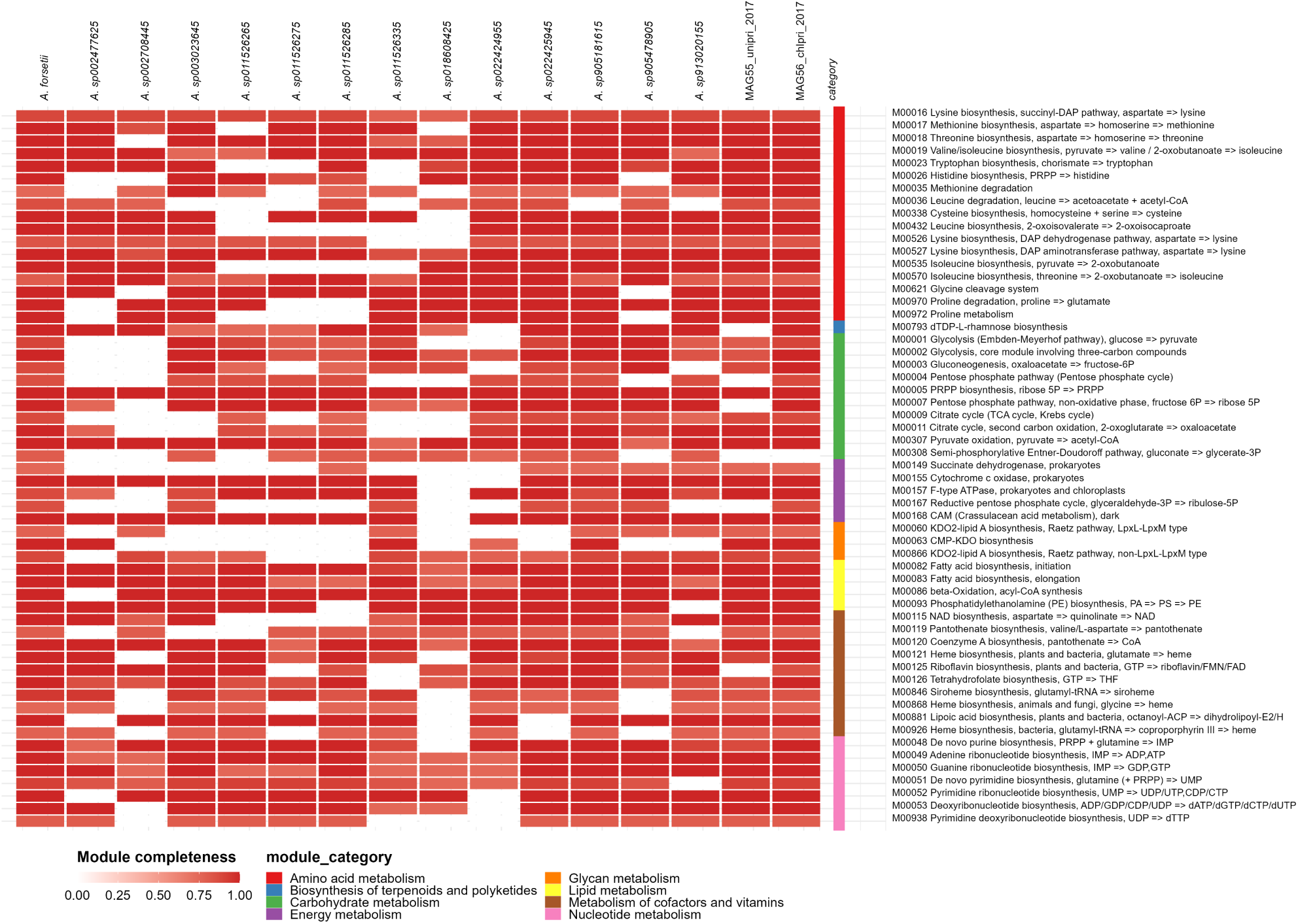
Completeness of KEGG metabolic modules in the genus ***Arcticimaribacter-2A* (order Flavobacteriales).** Two MAGs (55-56) were recovered in this genus, which relate to free-living seawater species. The color scale represents module completeness, from 0 to 1. Only modules with completeness ≥ 75% in at least one genome are presented.

**Suppl. Figure 25.**
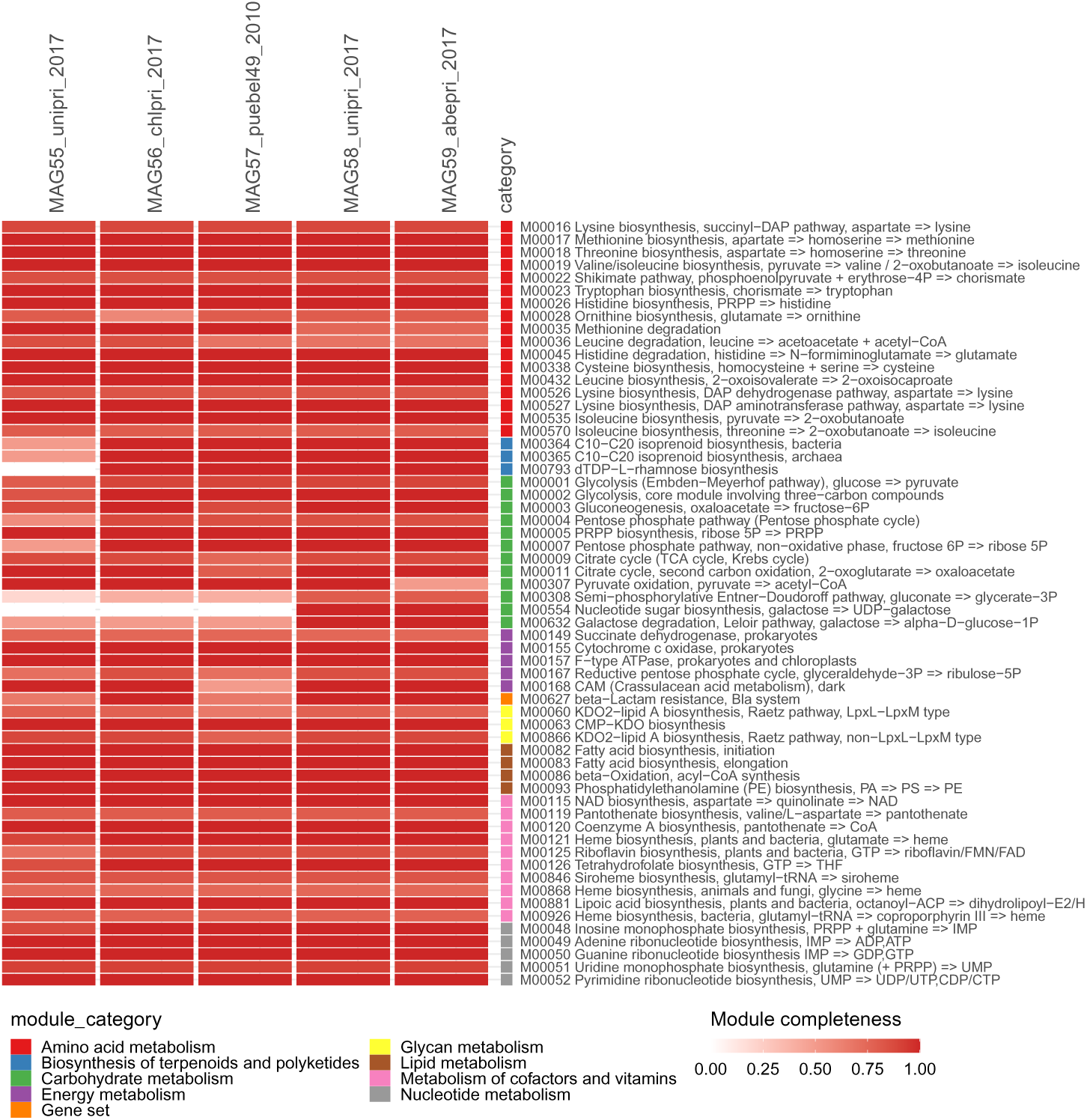
Completeness of KEGG metabolic modules for the five MAGs in the family Flavobacteriaceae. MAG55-MAG56 belong to genus *Arcticimaribacter* while MAG57-MAG59 belong to genus *MS024-2A*. The color scale represent the module completeness from 0 to 1. Only modules with completeness ≥ 75% in at least one genome are presented.

